# A conserved complex of microneme proteins mediates rhoptry discharge in *Toxoplasma*

**DOI:** 10.1101/2022.11.28.518173

**Authors:** Saima M. Sidik, Dylan Valleau, Yamilex Acevedo-Sánchez, Luiz C. Godoy, Charisse Flerida A. Pasaje, My-Hang Huynh, Vern B. Carruthers, Jacquin C. Niles, Sebastian Lourido

**Author notes:** Authors contributed equally to the work (Saima M. Sidik, Dylan Valleau).

## Abstract

Apicomplexan parasites discharge specialized organelles called rhoptries upon host cell contact to mediate invasion. The events that drive rhoptry discharge are poorly understood, yet essential to sustain the apicomplexan parasitic life cycle. Rhoptry discharge appears to depend on proteins secreted from another set of organelles called micronemes, which in *Toxoplasma gondii* includes MIC8 and the microneme-associated CRMP complex. Here, we examine the function of the microneme protein CLAMP, uncovering its essential role in rhoptry discharge. CLAMP forms a distinct complex with two other microneme proteins, the invasion-associated SPATR, and a previously uncharacterized protein we name CLAMP-linked invasion protein (CLIP). CLAMP-deficiency does not impact parasite adhesion or microneme protein secretion; however, knockdown of any member of the CLAMP complex affects rhoptry discharge. Phylogenetic analysis suggests orthologs of the essential complex components, CLAMP and CLIP, are ubiquitous across apicomplexans. Nevertheless, SPATR, which appears to act as an accessory factor in *Toxoplasma*, is essential during *Plasmodium falciparum* blood stages. Our results reveal a new protein complex that mediates rhoptry discharge following host-cell contact.

## INTRODUCTION

Apicomplexans are obligate unicellular parasites responsible for widespread diseases like malaria (*Plasmodium* spp.), cryptosporidiosis (*Cryptosporidium* spp.) and toxoplasmosis (*Toxoplasma gondii*). During the acute pathogenic stages of infection, apicomplexan parasites like *T. gondii* and *Plasmodium* spp. damage host cells through successive rounds of invasion, replication, and lysis (Bisio & Soldati-Favre, 2019; Blader *et al*, 2015). Active invasion of host cells by apicomplexans is mediated by specialized organelles—micronemes and rhoptries— that secrete proteins from the apical end of parasites in a highly regulated manner (Bisio & Soldati-Favre, 2019; Carruthers & Sibley, 1997). Secretion of the numerous small and oblong micronemes releases proteins throughout the extracellular stages of the parasite’s lytic cycle, from egress to invasion. By contrast, an arsenal of effector proteins is directly injected into host cells from the larger club-shaped rhoptries, in a process dependent on the prior secretion of micronemes (Kessler *et al*, 2008; Carruthers & Sibley, 1997). Rhoptry proteins locally reshape the host cell membrane in preparation for invasion and contribute to rewiring host cell signaling. Although essential for apicomplexan invasion, the regulatory cascade and proteinprotein interactions that mediate host cell detection and engage rhoptry secretion remain poorly understood.

Microneme proteins contribute to gliding motility, which supports displacement of parasites in tissues and invasion of host cells. Invasion additionally requires rhoptry discharge. Ca^2+^ signaling stimulates microneme fusion with the plasma membrane at the apical end of parasites to release proteins that perforate the host cell membrane, interact with the extracellular milieu, and bind host cell proteins (Carruthers & Sibley, 1999; Bisio & Soldati-Favre, 2019; Dubois & Soldati-Favre, 2019). Many microneme proteins remain tethered to the parasite’s plasma membrane following secretion and act as adhesins that allow parasites to bind host cells (Tomley & Soldati, 2001; Frénal *et al*, 2017). Some micronemal adhesins enable gliding motility by coupling the parasite’s actomyosin system to the host cell surface or extracellular matrix. Adhesion complexes are translocated rearward along the parasite surface to produce forward motility (Frénal *et al*, 2017). Some microneme proteins have dedicated roles during invasion; in *T. gondii*, the microneme protein MIC8 and the microneme-associated CRMP complex are required for rhoptry effector secretion, but not parasite motility (Sparvoli *et al*, 2022; Kessler *et al*, 2008; Singer *et al*, 2022). After secretion, some rhoptry effectors rewire host cell signaling to prevent innate immunity, while other effectors, notably the components of the rhoptry neck (RON) complex, prepare the site of rhoptry secretion for parasite invasion. A portion of RON2 becomes surface-exposed following translocation into the host cell, where it tightly binds the microneme protein AMA1. This AMA1 -RON2 complex mediates the tight apposition between host and parasite membranes at the site of invasion, called the moving junction, through which the parasite actively invades host cells using gliding-like motility (Frénal *et al*, 2017; Lamarque *et al*, 2014, 2011; Tyler & Boothroyd, 2011). The precise combination of surface-exposed microneme proteins and host cell molecules that triggers rhoptry discharge is not known.

Recent cryo electron tomography (cryo-ET) studies have begun to reveal the structure of the rhoptry secretion apparatus (RSA). The machinery involved in rhoptry discharge is more elaborate than the simple fusion of micronemes with the plasma membrane. Rhoptries docked to an apical vesicle that in turn binds the proteinaceous RSA centrally embedded in the apical plasma membrane (Mageswaran *et al*, 2021; Martinez *et al*, 2022; Aquilini *et al*, 2021). Correct assembly of the RSA complex is required for rhoptry discharge, as depletion of the protein Nd9 disrupts the RSA and blocks rhoptry protein secretion in *T. gondii* (Aquilini *et al*, 2021). Structural features of the RSA suggest that rhoptry proteins may be secreted through a channel at its center. Ca^2+^ signaling is also thought to contribute to rhoptry discharge. TgFER2, a conoid-associated Ca^2+^-binding protein, localizes to the cytosolic surface of the rhoptries and is essential for their discharge (Coleman *et al*, 2018). Moreover, treatment of parasites with Ca^2+^ ionophores leads to apparent fusion between the apical vesicle and the docked rhoptries (Segev-Zarko *et al*, 2022). The common reliance of both micronemes and rhoptries on Ca^2+^ signaling complicates deconvolution of the series of events that mediate invasion.

Proteins associated with rhoptry discharge appear conserved to different extents across the phylum. The RSA appears to be related to the secretory organelles of free-living ciliates, which are members of the superphylum Alveolata along with apicomplexans (Aquilini *et al*, 2021). Apicomplexans, therefore, appear to have adapted this conserved secretion machinery to their parasitic lifestyles. The CRMP proteins—required for rhoptry discharge but not the assembly of the apical vesicle or RSA—are broadly conserved among Alveolata much like the Nd proteins (Sparvoli *et al*, 2022) and bind additional microneme proteins in *T. gondii* (Singer *et al*, 2022). MIC8, a microneme protein required for rhoptry discharge in *T. gondii*, is limited to coccidians (Kessler *et al*, 2008) suggesting apicomplexans have clade-specific determinants of rhoptry discharge that potentially tune the process to their respective niches.

We previously demonstrated that the microneme protein CLAMP is necessary for *T. gondii* invasion of host cells (Sidik *et al*, 2016b). CLAMP is conserved across Apicomplexa, and its homolog is necessary for *P. falciparum* to progress through its lytic cycle, implying a conserved function. Here, we extend these results by identifying two additional microneme proteins that interact with CLAMP, and uncovering the essential function of this complex in rhoptry discharge. Our results strengthen the link between micronemes and rhoptries during invasion, adding to the understanding of the mechanisms that apicomplexan parasites use to establish their intracellular replicative niche.

## RESULTS

### CLAMP is a transmembrane microneme protein with a cytosolic C terminus

As previously reported, CLAMP is a transmembrane protein that colocalizes with the microneme marker MIC2 (**Fig. 1A**) (Sidik *et al*, 2016b). Several transmembrane microneme proteins have been investigated (Carruthers & Tomley, 2008; Dubois & Soldati-Favre, 2019), but none with the predicted topology of CLAMP. To examine the association of CLAMP with membranes, we prepared lysates from parasites in which the endogenous CLAMP locus was C-terminally tagged with the Ty epitope (CLAMP-Ty) (Bastin *et al*, 1996), and fractionated them by ultracentrifugation in the presence or absence of the detergent Triton X-100. Consistent with its membrane association, CLAMP co-fractionated with the GPI-anchored protein SAG1, moving from the pellet to the supernatant in the presence of detergent (**Fig. 1B**). By contrast, the cytosolic protein actin was largely found in the supernatant regardless of the presence of detergent.

**Figure 1.**
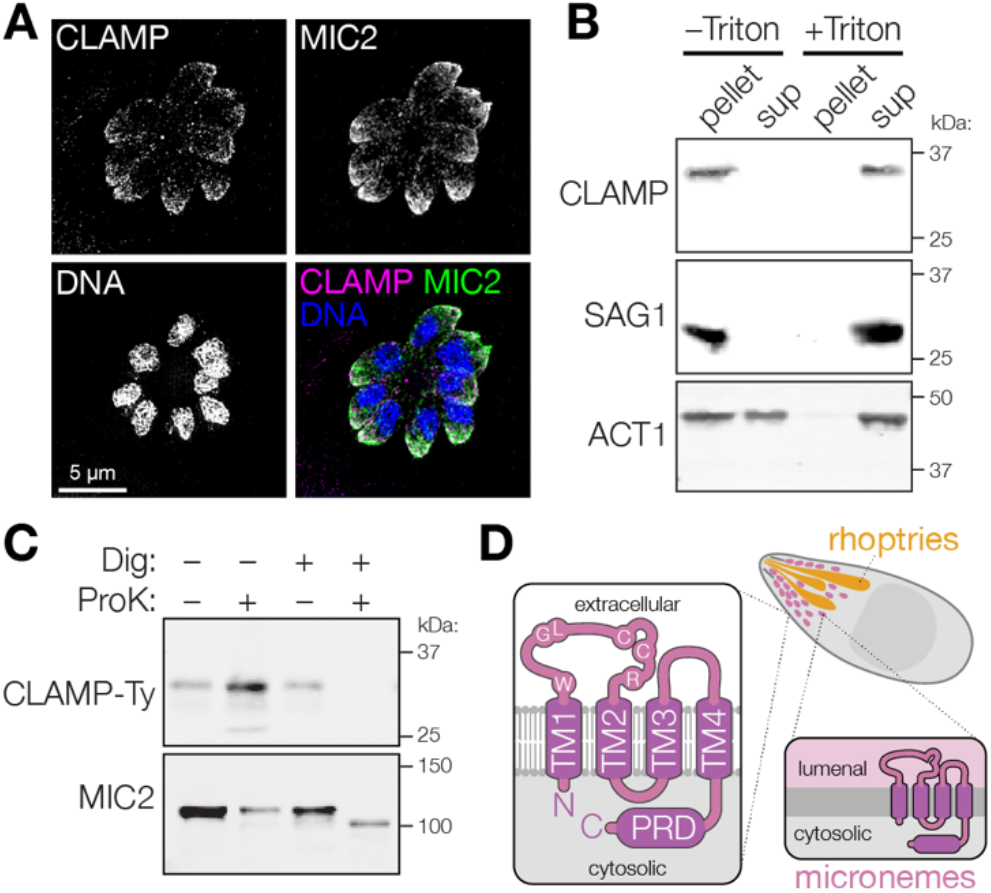
CLAMP is a microneme membrane protein with a cytosolic C terminus. **A**. Immunofluorescence microscopy revealed that CLAMP-Ty colocalizes with the microneme protein MIC2. **B**. Parasites expressing CLAMP-Ty were disrupted by freeze-thawing in the presence or absence of detergent (Triton X-100), then separated into pellet and supernatant (sup) fractions by ultracentrifugation. Anti-Ty immunoblotting revealed that CLAMP-Ty fractionates with the membrane protein SAG1, and differentially from cytosolic actin (ACT1). **C**. Sensitivity of CLAMP-Ty and MIC2 to Proteinase K digestion in intact parasites or those treated with digitonin to selectively disrupt their plasma membrane. Anti-Ty immunoblotting revealed that CLAMP’s cytosolic Ty tag was degraded in semi-permeabilized parasites while the luminal portion of MIC2 remained intact. **D**. Model of the topology of CLAMP in micronemes and on the plasma membrane following microneme exocytosis. Residues shared with claudins are highlighted.

Topology modeling (Dobson *et al*, 2015) predicts that CLAMP has four transmembrane helices (TMs), with its N and C termini extending into the cytosol. To confirm that CLAMP’s C terminus is cytosolic we used a Proteinase K protection assay in which the plasma membrane is gently permeabilized with digitonin to keep internal membranes intact and inaccessible to protease digestion. The single-pass transmembrane microneme protein MIC2 was used to calibrate the assay, based on availability of a monoclonal antibody against its lumenal domain and exposure of a short segment of its C terminus to the cytosol. Under conditions that preserved microneme integrity, the apparent molecular weight of MIC2 decreased when exposed to proteinase K, consistent with exclusive degradation of its cytosolic portion (**Fig. 1C**). Similarly, CLAMP’s cytosolic C-terminal Ty tag was degraded under conditions that preserved the lumenal MIC2 signal, in support of the predicted topology (**Fig. 1D**).

### CLAMP-deficient parasites establish normal contact with host cells prior to invasion

Our previous work showed that CLAMP-deficient parasites fail to invade host cells (Sidik *et al*, 2016b). Invasion is a multi-step process involving host cell attachment, secretion of microneme and rhoptry contents, and formation of the moving junction. To pinpoint the precise step of invasion affected by CLAMP knockdown, we generated a CLAMP conditional knockdown strain (CLAMP-HA cKD) using the U1 system. In this strain, a brief rapamycin pulse induces recombination of loxP sites flanking the 3^1^ UTR of the tagged gene and places an array of U1-binding sequences in line with the transcript, interfering with mRNA expression (Pieperhoff *et al*, 2015). We confirmed that CLAMP-HA cKD loses CLAMP expression when treated with rapamycin, as observed by immunoblot and immunofluorescence (**Fig. S1A–C**).

As many microneme proteins are thought to function as adhesins that facilitate host cell binding, we first quantified the ability of CLAMP-deficient parasites to adhere to human fibroblasts under flow conditions. Parasites were perfused over fibroblasts grown in microfluidic chambers, and we counted those that stably adhered to host cells for at least three seconds (**Fig. 2A**, **Supplemental Video 1**). In agreement with prior studies, pre-treating host cells with heparin inhibited parasite attachment (Monteiro *et al*, 1998). By contrast, CLAMP knockdown had no effect on *T. gondii* adhesion to host cells (**Supplemental Video 2**).

**Figure 2.**
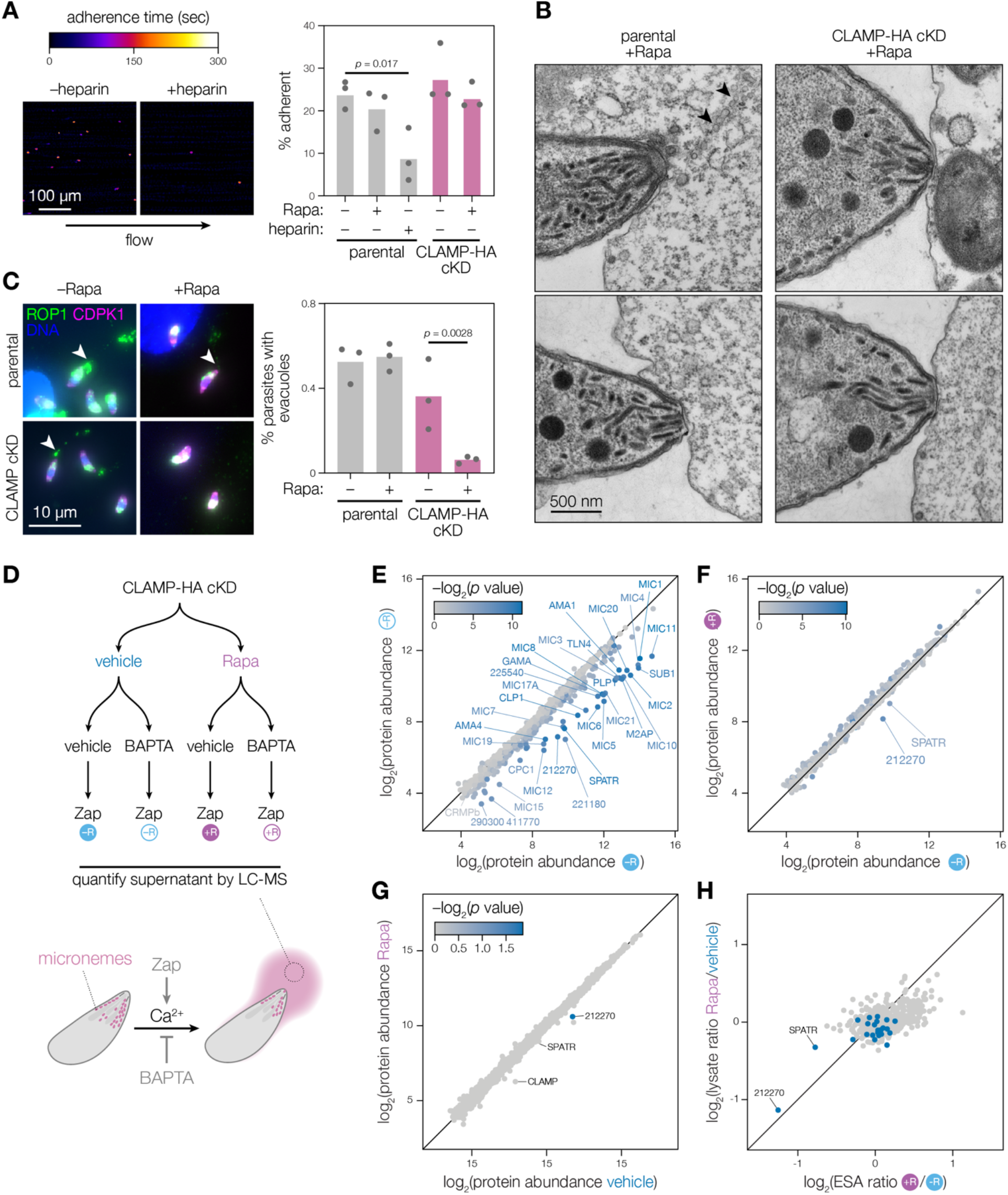
CLAMP loss specifically affects rhoptry discharge. **A**. Mycalolide B–treated parasites were imaged as they interacted with host cells under flow conditions. The percent of parasites remaining adhered to host cells for over 3 seconds was quantified. Pre-treating host cells with heparin reduced adhesion, but CLAMP-deficient parasites did not differ significantly from control strains. Mean is plotted for *n* = 3 biological replicates; *p* value for one-way ANOVA with Sidak’s multiple comparison test. **B**. Electron microscopy of mycalolide B–treated parasites interacting with K562 cells. Sections of the apical complexes of CLAMP cKD and parental strains treated with rapamycin (+Rapa) are shown. Black arrows indicate the position of structures reminiscent of evacuoles. **C**. Evacuole quantification of CLAMP cKD and parental parasites pre-treated with rapamycin (+Rapa) or vehicle (−Rapa). CLAMP depletion resulted in decreased association with evacuoles. Isolated parasites were pre-treated with cytochalasin D to prevent invasion before adding them to host cell monolayers. Evacuoles and parasites were visualized staining for ROP1 and CDPK1, respectively. Mean is plotted for *n* = 3 biological replicates; *p* value for one-way ANOVA with Sidak’s multiple comparisons test. **D**. Experimental setup for the quantitative mass spectrometric comparison of excretory-secretory antigens (MS-ESA). CLAMP cKD parasites were treated with zaprinast following pre-treatment with the calcium chelator BAPTA-AM or a vehicle control. **E**. Comparison of protein abundances in MS-ESA from the –Rapa samples treated with either vehicle (x axis) or BAPTA-AM (y axis) demonstrates detection of known secreted microneme proteins. Color scale indicates corrected *p* value for the comparison of the two axes. **F**. Comparison of MS-ESA abundance from induced, zaprinast-treated samples comparing the CLAMP cKD following pre-treatment with vehicle (x axis; as in E) or rapamycin (y axis). **G**. Quantifying the abundance of proteins in the total lysate of parasites in either the −Rapa or +Rapa conditions used for the MS-ESA. CLAMP depletion affects the abundance of only certain proteins. **H**. Comparison of the relative ratios of secretion to the overall abundance when CLAMP is present or knocked down shows that TGGT1_212270 abundance decreases in both the ESA fraction and the parasite lysate when CLAMP is depleted, while SPATR secretion is more affected than its overall protein abundance. The microneme proteins annotated in E are displayed in blue.

Following adhesion, parasites reorient to make apical contact with the host cell plasma membrane. To examine the ultrastructure of the parasite apical end during this stage of the invasion process, we performed transmission electron microscopy on the CLAMP cKD and its parental strain as they interacted with K562 human lymphoblast cells. The non-adherent nature of K562 cells facilitated preparations for electron microscopy. Parasites were pre-treated with mycalolide B to prevent actin-based invasion without affecting host cell adhesion (Cirelli *et al*, 2014). No morphological differences could be observed between the apical ends of the CLAMP-HA cKD and its parental strain, irrespective of rapamycin treatment (**Fig. S2**). In both cases, examples of tightly apposed parasite apical ends and regions of the host plasma membrane could be observed, regardless of CLAMP knockdown (**Fig. 2B**). In rare cases, apical ends from the parental strain were associated with trails of small vacuoles, reminiscent of the previously described evacuoles that result from rhoptry discharge (Håkansson *et al*, 2001). Similar structures were not observed in proximity to CLAMP-knockdown parasites. Nevertheless, these results demonstrate that, in the absence of CLAMP, parasites retain the ability to correctly reorient and establish apical contact with host cells, despite failing to initiate invasion.

### CLAMP is required for rhoptry discharge and secretion of two other microneme proteins

Following microneme-dependent apical attachment to host cells, parasites discharge rhoptries. Rhoptry discharge is essential, as the secreted proteins include critical components of the tight junction required for invasion of host cells (Carruthers & Sibley, 1997; Ben Chaabene *et al*, 2021). Inhibition of parasite gliding motility (e.g., preventing actin polymerization) blocks invasion but does not affect rhoptry discharge, which can be observed by examining the localization of rhoptry proteins to evacuoles (Håkansson *et al*, 2001; Sibley, 2010; Kessler *et al*, 2008). To determine whether CLAMP-deficient parasites secrete rhoptry proteins, we quantified evacuole formation. CLAMP-depleted parasites and controls were pre-treated with the actin polymerization inhibitor cytochalasin D and allowed to interact with host cells for a few minutes prior to fixation. Staining for the rhoptry protein ROP1 revealed that 30–50% of control parasites were associated with evacuoles (**Fig. 2C**, arrows). By contrast, evacuoles were only associated with 6% of CLAMP-deficient parasites—low enough to be attributable to the small proportion of CLAMP-HA cKD parasites that fail to undergo recombination. We conclude that loss of CLAMP results in a failure to discharge rhoptries, preventing invasion.

Only two groups of surface-exposed proteins, MIC8 and the CRMPs, have been specifically associated with rhoptry discharge in *T. gondii* (Kessler *et al*, 2008; Singer *et al*, 2022; Sparvoli *et al*, 2022). We previously showed that CLAMP depletion does not affect secretion of the microneme protein MIC2 (Sidik *et al*, 2016b). Nevertheless, the similarity between the CLAMP phenotype and that of the CRMPs and MIC8 motivated a systematic evaluation of CLAMP’s role in microneme secretion. We developed a quantitative mass spectrometry–based method to measure the excretory-secretory antigens of *T. gondii* strains (MS-ESA; **Fig. 2D**). Zaprinast is a phosphodiesterase inhibitor that triggers Ca^2+^ release from intracellular stores by activating the cGMP-dependent protein kinase (Yuasa *et al*, 2005; Lourido *et al*, 2012; Sidik *et al*, 2016a; Brown *et al*, 2016). Since micronemes are secreted in a Ca^2+^-dependent manner, we first compared the ESA fractions from zaprinast-stimulated parasites that had been pre-treated with either the Ca^2+^-chelator BAPTA-AM or a vehicle control (Carruthers & Sibley, 1999). As anticipated, microneme proteins were more abundant in the ESA from the vehicle control compared to the BAPTA-AM-treated parasites (**Fig. 2E**). 27 of the proteins significantly enriched from parasites with an intact Ca^2+^-signaling pathway were either annotated as microneme proteins or found to be microneme-localized by spatial proteomics (Barylyuk *et al*, 2020) (**Fig. 2E**). Our results demonstrate that the MS-ESA approach is a valid way of examining *T. gondii* microneme secretion.

Our MS-ESA results highlight several potential microneme proteins that have been poorly characterized or only recently identified. Among the Ca^2+^-dependent secreted proteins we found MIC19—which was not predicted to be microneme localized by spatial proteomics—as well as the recently identified microneme protein MIC21 (Tagoe *et al*, 2021; Barylyuk *et al*, 2020). We also found CRMPb, which inconsistently localizes with micronemes and forms a complex with CRMPa and MIC15 (Singer *et al*, 2022). Several additional proteins secreted in a Ca^2+^-dependent manner have not been examined in detail. TGGT1_212270 and TGGT1_221180, were annotated as microneme-localized by spatial proteomics, but have not been further studied (Barylyuk *et al*, 2020). TGGT1_411770 has not been localized by other datasets, but its paralog TGGT1_319090 is predicted to be microneme-localized. It is possible that some of the proteins secreted in a Ca^2+^-dependent manner do not originate from micronemes. TGGT1_225540 and TGGT1_290300 were also secreted in a Ca^2+^-dependent manner yet were not microneme-localized by spatial proteomics.

Using MS-ESA, we studied the effect of CLAMP knockdown on microneme secretion. Nearly all microneme proteins were secreted to a similar extent in the presence or absence of CLAMP (**Fig. 2F**). Notably, MIC8 and CRMP complex components were unaffected by CLAMP repression. However, two proteins showed a striking CLAMP dependency: SPATR and one of the previously unstudied proteins, TGGT1_212270. We verified whether changes in SPATR and TGGT1_212270 abundance in the ESA samples were due to a secretion defect or protein destabilization. Measuring the relative abundance of proteins in the lysates of rapamycin or vehicle-treated CLAMP-HA cKD by quantitative mass spectrometry, the only significantly depleted protein was TGGT1_212270 (**Fig. 2G**). Comparing the abundance ratios of proteins in the ESA fraction and whole-parasite lysates suggests that CLAMP knockdown leads to decreased TGGT1_212270 stability, whereas the reduction of SPATR in the ESA fraction is likely due to a secretion or trafficking defect (**Fig. 2H**). Prior studies suggest that SPATR and TGGT1_212270 localize to micronemes where CLAMP could influence their trafficking and stability (Barylyuk *et al*, 2020; Sidik *et al*, 2016b; Huynh *et al*, 2014).

### CLAMP, CLIP, and SPATR comprise a stable protein complex

We used co-immunoprecipitation to identify proteins that interact with CLAMP. Using parasites in which CLAMP was endogenously tagged with the Ty epitope (CLAMP-Ty), we immunoprecipitated CLAMP-Ty from detergent-solubilized parasite lysates (**Fig. 3A**), and analyzed captured proteins by mass spectrometry. Throughout our experiments, CLAMP has been poorly detected by MS, despite being clearly observed in the samples by immunobloT. We attribute its absence from our MS results to its hydrophobicity and unusual pattern of tryptic fragments. Nevertheless, comparing the fold enrichment in protein abundance between immunoprecipitations from CLAMP-Ty parasites over their parental counterparts revealed two proteins enriched in both biological replicates: SPATR and TGGT1_212270 (**Fig 3B**)—notably, the two CLAMP-dependent constituents in the ESA (**Fig. 2F**).

**Figure 3.**
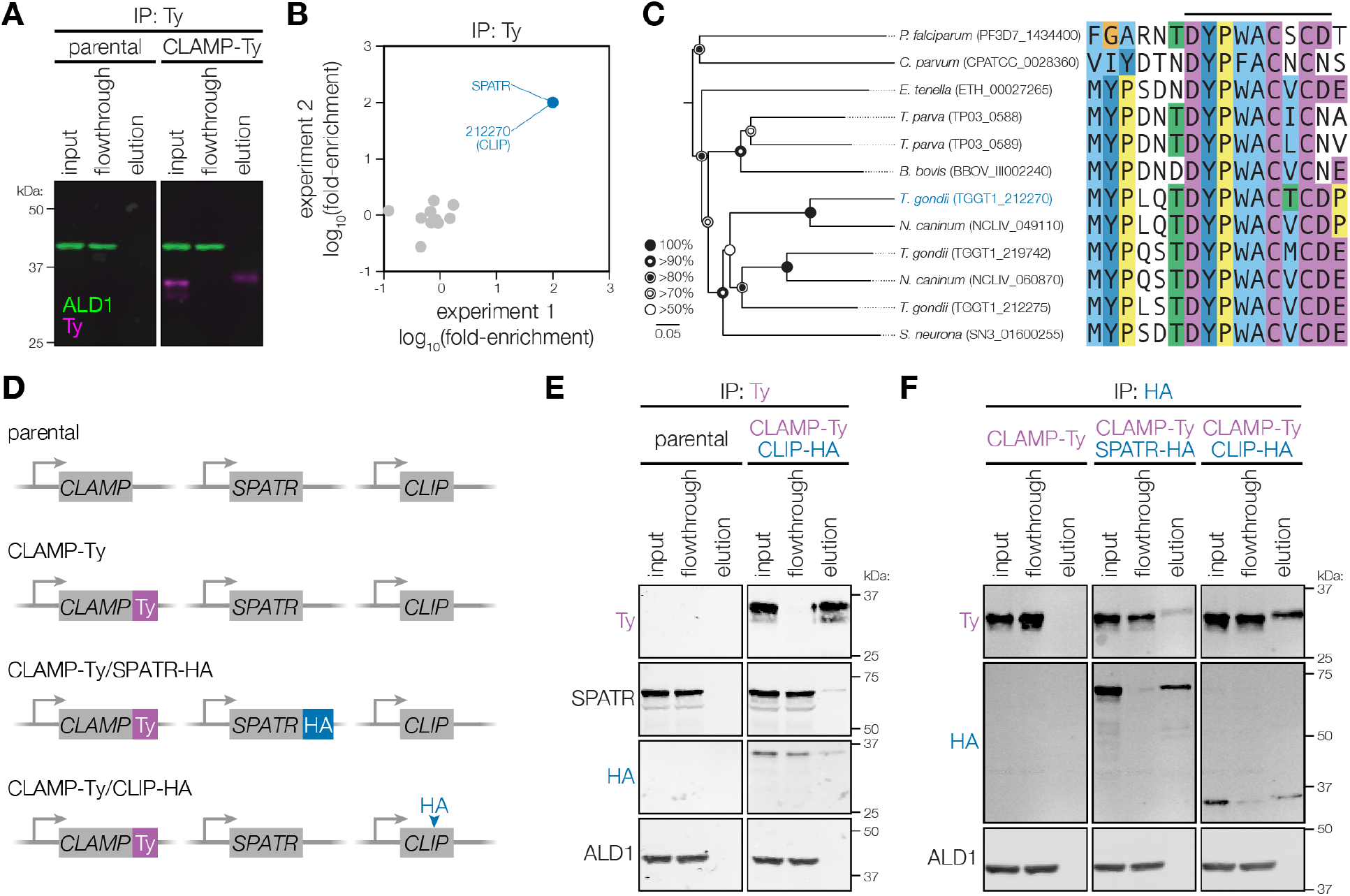
CLAMP forms a complex with CLIP (TGGT1_212270) and SPATR. **A**. Immunoblot of samples from an immunoprecipitation of CLAMP-Ty comparing the tagged strain to the parental control. ALD1 is used as a soluble antigen to monitor the specificity of the enrichment. **B**. SPATR and TGGT1_212270 (CLIP) were reproducibly co-immunoprecipitated with CLAMP-Ty compared to the untagged control. Label-free quantification of eluates by mass spectrometry was used to calculate fold-enrichment. **C**. Neighbor-joining phylogenetic tree of selected CLIP homologs from across the apicomplexan phylum. The region around conserved motif 1 is shown within the sequence alignment. Bootstrap values for 1,000 trials are displayed. Abbreviated species names are provided for *Toxoplasma gondii, Plasmodium falciparum, Theileria parva, Babesia bovis, Neospora caninum, Sarcocystis neurona, Hammondia hammondi*, *Eimeria tenella*, and *Cryptosporidium parvum*. An extended tree and additional conserved motifs are provided in Fig. S3. **D**. Diagram of strains carrying HA tags on CLIP and/or SPATR. SPATR was tagged at its C terminus and CLIP internally, following Arg210. **E– F**. Immunoblot of samples from an immunoprecipitation of CLAMP-Ty (E) or the HA-tagged interacting partners (F) demonstrating the interaction of all three proteins.

Based on the observed protein-protein interaction, we named TGGT1_212270 CLAMP-Linked Invasion Protein (CLIP). Searches of apicomplexan sequences (veupathdb.org) using the basic local alignment search tool (BLAST) identified multiple CLIP-related sequences broadly distributed across the apicomplexan phylum. Examining the sequences revealed a highly conserved motif near the C-terminal TM of homologous sequences (DYP[WF]ACxC), which recovered 1–3 matching sequences from all apicomplexan genomes examined (**Fig. 3C**, and **Fig. S3**). Most apicomplexans have a single ortholog, including malaria-causing parasites from the genus *Plasmodium*. However, gene-duplication appears to have given rise to groups of paralogs, with *Theileria* spp. harboring two, and most cyst-forming coccidians like *T. gondii* harboring three. The early branching coccidian *Sarcocystis neurona* appears to have a single CLIP ortholog. Both CLIP paralogs in *T. gondii* appear to be dispensable during the tachyzoite stages (Sidik *et al*, 2016b) and show peaks of expression during the chronic stages (TGGT1_212275) or in the definitive host (TGGT1_219742) (Ramakrishnan *et al*, 2019), suggesting they might function during other stages of the life cycle. MEME analysis (Bailey & Elkan, 1994; Bailey *et al*, 2009) of the CLIP-related sequences revealed additional regions of high conservation (**Fig. S3**), despite overall low sequence similarity. We conclude that CLIP may have as broad a pattern of conservation as previously suggested for CLAMP (Sidik *et al*, 2016b), but appears to be under stronger positive selection.

We generated additional strains to validate the putative interactions between CLAMP, CLIP, and SPATR. Using CRISPR-mediated homologous recombination, we generated a strain in which SPATR was C-terminally HA tagged in the CLAMP-Ty background (CLAMP-Ty/SPATR-HA); however, attempts to modify the *CLIP* locus for N- or C-terminal tagging proved unsuccessful. Based on the possible presence of N-terminal secretion signals and the high degree of conservation along the C terminus of the protein, we successfully targeted a less conserved internal region, after Arg210 of the coding sequence, for marker-free integration of an HA epitope (CLAMP-Ty/CLIP-HA; **Fig. 3D**). We immunoprecipitated CLAMP-Ty and used immunoblotting to confirm its interaction with both CLIP and SPATR (**Fig. 3E**). We also performed reciprocal immunoprecipitations using the HA-tags of the interacting partners and found that CLIP-HA and SPATR-HA both co-immunoprecipitated CLAMP (**Fig. 3F**). These experiments provide strong evidence for a stable three-way complex between CLAMP, CLIP, and SPATR.

### CLIP is an essential secreted microneme protein

We sought to confirm the subcellular localization of CLIP. Using super-resolution microscopy on the strain in which CLIP was HA tagged, we found that, like CLAMP (**Fig. 1A**) and SPATR (Huynh *et al*, 2014), CLIP adopts a punctate apical pattern that largely colocalizes with the microneme marker MIC2 (**Fig. 4A**). Differential ultracentrifugation of parasite lysates in the presence or absence of detergent confirmed the membrane association of CLIP, based on its co-fractionation with the known membrane protein SAG1 (**Fig. 4B**). This is consistent with an *in silico* topology analysis (Dobson *et al*, 2015) predicting that CLIP has a single transmembrane helix at its C terminus. CLIP peptides are detected in the Ca^2+^-dependent ESA fraction, consistent with microneme secretion (**Fig. 2E**). These results suggest the N terminus of CLIP largely resides in the microneme lumen or is surface exposed following microneme exocytosis (**Fig. 4C**).

**Figure 4.**
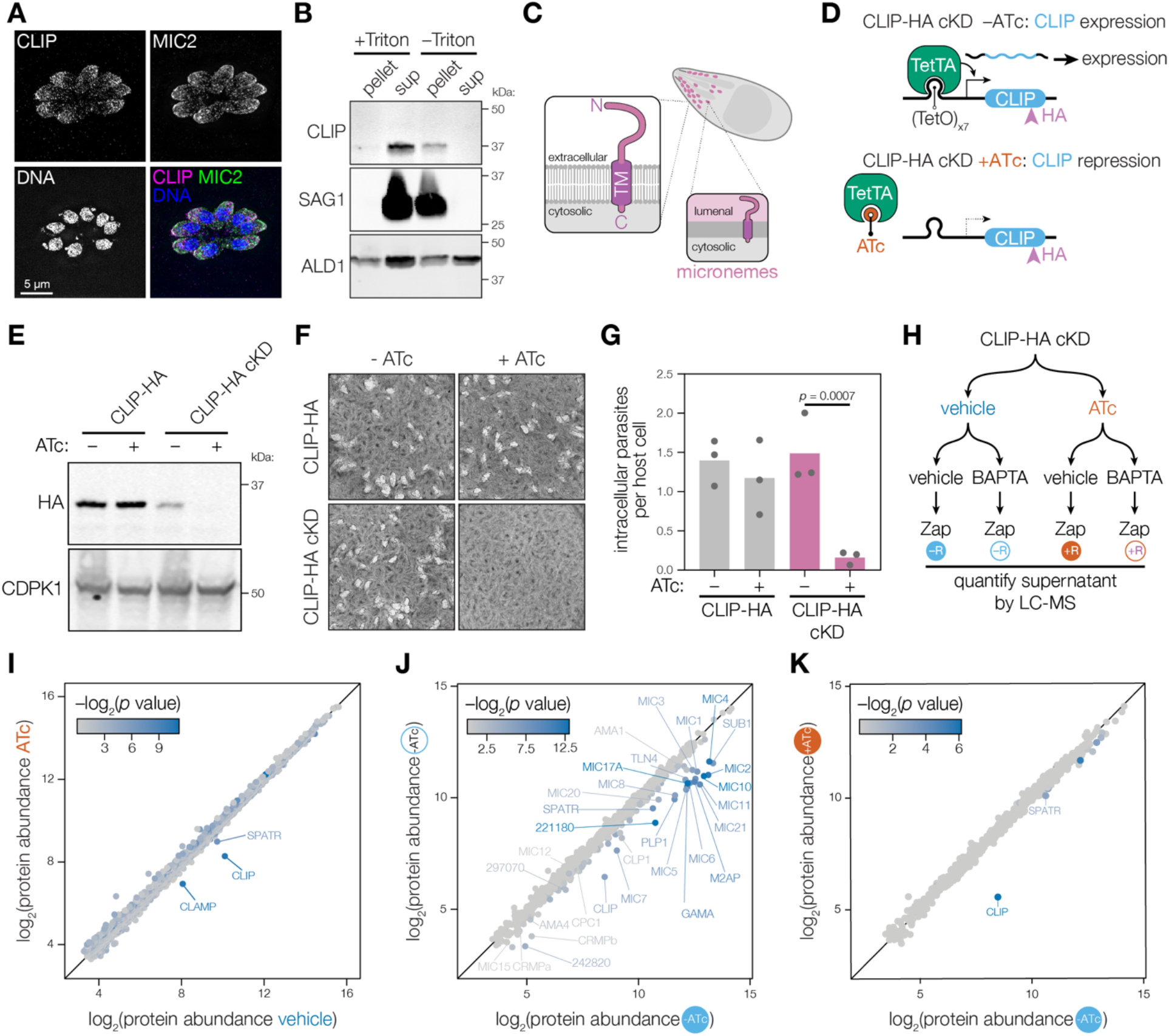
CLIP is a secreted single-pass transmembrane microneme protein that is necessary for invasion. **A**. Immunofluorescence staining with anti-HA reveals that CLIP-HA colocalizes with the microneme marker MIC2. **B**. CLAMP-Ty/CLIP-HA parasites were disrupted using multiple freeze/thaw cycles, then fractionated by ultracentrifugation in the presence or absence of detergent (Triton X-100). CLIP partitions similarly to the membrane protein SAG1 and differently from the cytosolic protein Aldolase. **C**. Our data support a model in which CLIP is a single-pass transmembrane microneme protein that is secreted onto the parasite’s surface, then shed into the extracellular space with an exposed ectodomain. **D**. The CLIP conditional knockdown strain (CLIP cKD) was made by replacing the region 5’ to the coding sequence of CLIP-HA with the *TetO7-SAG1* promoter in a Tet-transactivator expressing background (TetTA). Addition of anhydrotetracycline (ATc) results in release of the TetTA and inhibition of transcription. **E**. Growth in the presence of ATc for 48 h leads to conditional knockdown of CLIP-HA cKD as measured by anti-HA immunoblot. CDPK1 is used as a loading control. **F**. CLIP is essential to the lytic cycle. CLIP-HA cKD parasites failed to form plaques when treated with ATc, while the CLIP-HA parental strain formed plaques regardless of ATc treatment. **G**. CLIP-HA cKD parasites fail to invade host cells after 48 h of ATc treatment. Mean is plotted for *n* = 3 biological replicates; *p* values derive from a one-way ANOVA followed by Sidak’s multiple comparison test. **H**. Experimental setup for the MS-ESA and whole proteome analysis of CLIP conditional knockdown. CLIP-HA cKD parasites that had CLIP knocked down with ATc or were treated with vehicle control for 48 h. For the MS-ESA, parasites were then stimulated with zaprinast following pre-treatment with the Ca^2+^ chelator BAPTA-AM or a vehicle control. **I–K**. Depletion of CLIP by 48 h ATc treatment (y axis) did not dramatically affect the stability of proteins other than CLIP in total parasite lysate compared to vehicle control (x axis; I). Microneme proteins are generally found in the ESA fraction of the vehicle-treated CLIP-HA cKD strain as shown by treatment with BAPTA-AM to inhibit secretion (y axis) or a vehicle control (x axis; J). Comparison of the MS-ESA for CLIP knockdown (+ATc, y axis) to vehicle control (−ATc, x axis) reveals that CLIP knockdown leads to depletion of CLIP, CLAMP, and SPATR in the ESA fraction (K). Color scale indicates corrected *p* value for the comparison of the two axes for I-K.

Both CLAMP and SPATR mutants have invasion defects (Huynh *et al*, 2014; Sidik *et al*, 2016b), which in the case of CLAMP, leads to a complete block in the parasite’s lytic cycle. To determine whether CLIP knockdown phenocopies that of its binding partners, we constructed an inducible CLIP-HA cKD mutant using the Tet-off transactivator system (Meissner *et al*, 2002) (**Fig. 4D**). We confirmed that growth of CLIP-HA cKD in anhydrotetracycline (ATc) leads to loss of CLIP-HA expression as observed by immunoblot and immunofluorescence assays (**Fig. 4E**, **Fig. S4A**). CLIP-HA cKD parasites also failed to form plaques in the presence of ATc (**Fig. 4F**). These results demonstrate that CLIP is essential for completion of the parasite lytic cycle, consistent with its low phenotype score (−3.05) in our genome-wide screens (Sidik *et al*, 2016b). Invasion assays showed that this lytic cycle arrest is likely due to an inability of CLIP xknockdown parasites to enter host cells (**Fig. 4G**), as is the case with CLAMP knockdown.

To assess whether CLIP affects secretion or stability of additional microneme proteins, as observed for CLAMP, we performed whole-proteome profiling and an MS-ESA assay of the CLIP-HA cKD strain (**Fig. 4H**). CLIP depletion did not lead to a significant destabilization of any other proteins in parasite lysate (**Fig. 4I**). As with the CLAMP MS-ESA, microneme proteins were found in the calcium-dependent ESA fraction of zaprinast-induced CLIP-HA cKD confirming intact microneme secretion in the absence of CLIP (**Fig. 4J**). During CLIP knockdown, the protein most dramatically depleted from the ESA fraction was CLIP itself, with a modest effect on SPATR secretion (**Fig. 4K**); however, CLAMP was not detected in this experiment.

Many single-pass transmembrane microneme proteins are released from the plasma membrane by parasite-encoded rhomboid proteases (Dowse *et al*, 2005; Opitz *et al*, 2002). However, CLIP lacks a typical rhomboid protease cleavage site (Sheiner *et al*, 2010), raising the possibility that the extracellular domain is cleaved outside of the predicted C-terminal transmembrane helix and N-terminal to the internal HA tag of CLIP-HA parasites. We therefore generated a rabbit anti-CLIP polyclonal antibody against the N-terminal region of CLIP, comprising residues 1–187 (**Fig. S4B**). We immunoblotted the ESA fractions from our MS-ESA assay, finding two major ~20 kDa bands detected by the anti-CLIP-N antibody in the ESA fraction. The CLIP bands were lost when CLIP-HA was depleted by ATc treatment (**Fig. S4C**). These results suggest that the N-terminal ectodomain of CLIP is released from the parasite plasma membrane into the supernatant. We conclude that CLAMP and CLIP most likely form a membrane-embedded complex, with the CLIP ectodomain, CLAMP loops 1 and 3, and SPATR exposed on the parasite surface after secretion from micronemes.

### The CLAMP complex is necessary for rhoptry secretion

We examined whether, like CLAMP, CLIP and SPATR are also necessary for rhoptry secretion. Because the morphology of evacuoles is extremely variable, counting these structures is difficult to automate and subject to investigator bias. Others have measured rhoptry secretion by assessing the phosphorylation of the host transcription factor STAT6, which is targeted by the rhoptry kinase ROP16 following secretion into host cells (Ong *et al*, 2010; Butcher *et al*, 2011; Saeij *et al*, 2006; Lamarque *et al*, 2014). We incubated mycalolide B–treated parasites with human fibroblasts, which we then stained for host phospho STAT6 (pSTAT6). Using a custom analysis script, we quantified the percentage of host cells with nuclear pSTAT6 (**Fig. 5A**) as an unbiased readout for rhoptry secretion.

**Figure 5.**
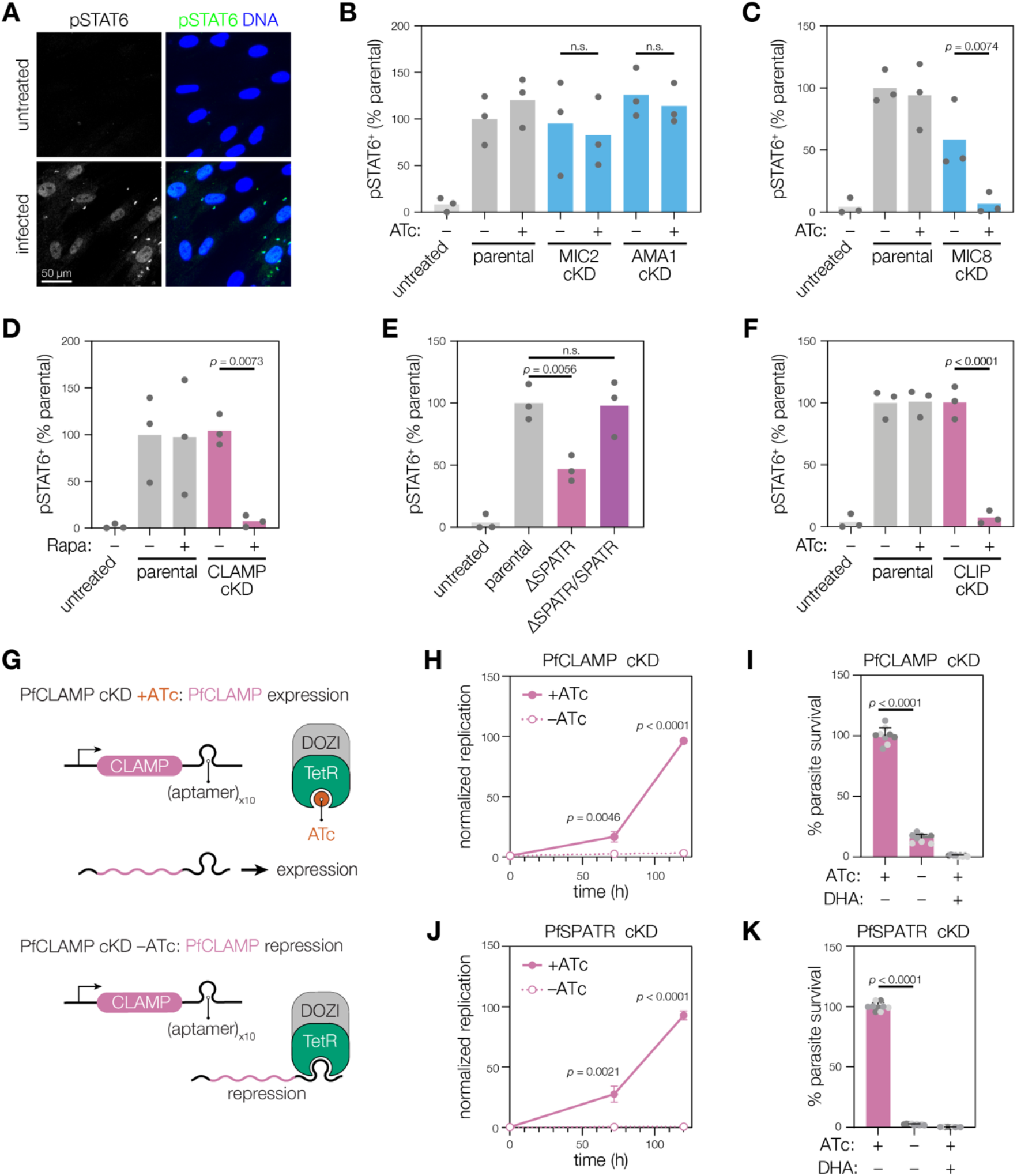
All members of the CLAMP complex are necessary for robust secretion of rhoptry contents. **A**. Examples of HFF cells stained with anti-pSTAT6 and Hoechst (to visualize DNA) after treatment with mycalolide B-treated parasites. **B–C**. Anhydrotetracycline (ATc) treatment did not significantly change the ability of MIC2-cKD or AMA1 -cKD parasites to elicit pSTAT6-positive host cell nuclei (B). However, ATc treatment significantly reduced the ability of MIC8-cKD parasites to elicit pSTAT6-positive host cell nuclei (C). Mean is plotted for *n* = 3 biological replicates; *p* values derive from one-way ANOVA followed by Sidak’s multiple comparison test prior to data normalization. **D–F**. All members of the CLAMP complex are required for robust nuclear translocation of host pSTAT6. Mean is plotted for *n* = 3 biological replicates; *p* values derive from one-way ANOVAs followed by Sidak’s multiple comparison tests prior to data normalization. **G**. The *P. falciparum* conditional knockdown system used to downregulate CLAMP or SPATR. Removal of ATc from the culture medium allows transcript binding of the TetR-DOZI fusion protein to the Tet aptamer sequences and leads to conditional loss of protein translation. **H–K**. Conditional knockdown of *Pf*CLAMP (H–I) or *Pf*SPATR (J–K) by removal of ATc from the culture medium leads to complete loss of parasite growth. Activity of constitutively expressed *Renilla* luciferase was used to track growth of each *P. falciparum* conditional knockdown line. Dihydroartemisinin (DHA) is used as a control for growth inhibition (I and K); *p* values derive from multiple *t*-tests for each time point using the Benjamini-Hochberg FDR approach (H and J) or one-way ANOVA followed by Sidak’s multiple comparison test (I and K).

We benchmarked the pSTAT6 assay using conditional mutants for various microneme proteins. The adhesin MIC2 mediates attachment to host cells during gliding motility and the moving junction component AMA1 forms stable interactions during invasion, but neither is expected to influence rhoptry discharge (Huynh & Carruthers, 2006; Lamarque *et al*, 2014). As expected, knocking down MIC2 or AMA1 did not substantially alter the number of host cells displaying nuclear accumulation of pSTAT6 (**Fig. 5B**). On the other hand, MIC8 has been shown to be necessary for rhoptry secretion (Kessler *et al*, 2008) and its knockdown resulted in a near complete block in rhoptry discharge (**Fig. 5C**). Based on these results we conclude that our assay robustly distinguishes between effects on rhoptry exocytosis and other mechanisms that impact invasion.

We next examined the effect of depleting different components of the CLAMP complex on rhoptry discharge. As expected from quantification of evacuoles, knockdown of CLAMP severely diminished the number of phospho-STAT6 positive nuclei observed in infected cultures (**Fig. 5D**). To examine the role of SPATR, we took advantage of strains in which *SPATR* was replaced with a chloramphenicol acetyltransferase cassette (*ΔSPATR*) and later complemented (*ΔSPATR/SPATR*) (Huynh *et al*, 2014). Loss of SPATR reduced the number of host cells that accumulate pSTAT6 by approximately one half (**Fig. 5E**), consistent with the modest invasion defect reported for the mutant (Huynh *et al*, 2014) and its minimal impact on fitness in genome-wide screens (Sidik *et al*, 2016b). During CLIP knockdown, we observed a near complete loss of pSTAT6 in host cell nuclei, replicating the phenotype seen with CLAMP knockdown (**Fig. 5F**). We conclude that the microneme protein complex composed of CLAMP, CLIP, and SPATR is necessary for secretion of rhoptry contents.

### CLAMP and SPATR are essential for blood-stage growth of *Plasmodium falciparum*

All three members of the CLAMP complex have homologs throughout the Apicomplexa, including in the causative agent of the most severe forms of malaria, *Plasmodium falciparum*. Though *Pf*CLIP has been resistant to N or C terminal tagging, we were able to construct conditional knockdown strains for both *Pf*CLAMP and *Pf*SPATR by inserting tandem Tet repressor (TetR) aptamer sequences at the endogenous locus in a parental strain expressing the Tet repressor fused to the translational repressor DOZI (TetR-DOZI), as well as constitutively expressing *Renilla* luciferase. When ATc is removed from the culture media, TetR-DOZI can bind the transcript and inhibit protein translation (**Fig. 5G**). Using *Renilla* luciferase activity as a metric for growth, we confirmed that *Pf*CLAMP is essential for bloodstage growth of *P. falciparum* (Sidik *et al*, 2016b). When *Pf*SPATR expression was repressed by the removal of ATc, we found a similarly complete block of blood-stage growth, demonstrating that *Pf*SPATR and *Pf*CLAMP are both essential during blood stage growth of *P. falciparum*.

## DISCUSSION

The microneme protein CLAMP was previously identified as an invasion factor necessary for *T. gondii* and *P. falciparum* fitness (Sidik *et al*, 2016b). Here, we discover that CLAMP is part of a complex together with two additional microneme proteins: SPATR and the newly characterized protein CLIP (TGGT1_212270). All members of this complex are necessary for normal secretion of rhoptry contents—with the membrane-associated CLIP and CLAMP being absolutely required. CLAMP knockdown parasites are unaltered in host cell adhesion, microneme secretion, or apical complex morphology, leading us to conclude that their invasion defect stems specifically from an inability to discharge rhoptries.

Microneme proteins perform diverse functions as parasites spread between host cells, including membrane disruption during egress, adhesion during gliding motility, and establishment of the moving junction during invasion. However, only a handful of microneme proteins appear to mediate rhoptry discharge. MIC8, a prototypical single-pass transmembrane microneme protein, was the first factor shown to be specifically required for rhoptry discharge in *T. gondii* (Kessler *et al*, 2008). More recently, two large multipass transmembrane proteins (CRMPa and CRMPb) were similarly associated with rhoptry discharge and shown to form a complex with two single-pass microneme proteins bearing thrombospondin domains (Singer *et al*, 2022; Sparvoli *et al*, 2022). Although the CRMP complex proteins do not localize to micronemes unambiguously (Singer *et al*, 2022; Barylyuk *et al*, 2020), they accumulated at the apical end of extracellular parasites (Sparvoli *et al*, 2022). Despite extensive analysis through co-immunoprecipitation, there is no evidence that MIC8 and the CRMP complex interact with one another or with the CLAMP complex, suggesting the three independently contribute to rhoptry discharge. The two complexes and MIC8 possess domains that could potentially interact with the surface of host cells (Kessler *et al*, 2008; Huynh *et al*, 2014; Sparvoli *et al*, 2022), implying that coordinated binding to various host cell molecules may determine the site of apical adhesion and rhoptry discharge.

The mechanics of how microneme proteins induce rhoptry content secretion have not been established. Unlike microneme or dense granule discharge, the release of rhoptry contents occurs specifically upon contact with host cells. Rhoptry discharge therefore requires transduction of molecular or physical cues that signal attachment to host cells. CLAMP was originally named for its similarity to claudins although its C terminus is distinct, possessing a proline-rich domain (PRD) that extends into the parasite cytosol in place of the PDZ-binding motif that mediates assembly of tight junctions by most mammalian claudins (Hou *et al*, 2013). No function has been attributed to the PRD, but its orientation may enable the CLAMP complex to interact with the RSA or signaling pathways following the apical attachment of parasites with the host plasma membrane. Our data indicate that apical attachment to host cells is indeed intact in the absence of CLAMP, placing the function of the CLAMP complex downstream from general physical attachment to host cells.

The relationship between the CLAMP complex and the RSA remains unclear. The specific identities of proteins forming the RSA have not been confirmed; however, knockdown of the non-discharge protein Nd9 abolishes proper RSA assembly (Mageswaran *et al*, 2021). The CLAMP complex does not co-precipitate with the Nd9 interactome in our experiments, nor in prior work (Aquilini *et al*, 2021), suggesting that CLAMP does not reside at the RSA, at least in absence of host cell apposition. By contrast, components of the CRMP complex do colocalize with the RSA prior to invasion, but dissociate from the apical end of the parasite once the moving junction is formed (Sparvoli *et al*, 2022). The transient association of the CRMP complex with the RSA led to speculation that it serves to detect and transduce the signals of host-cell attachment, as we propose for the CLAMP complex. In the complex milieu of animal tissues, distinguishing attachment to host cells from adherence to extracellular matrix components may require the use of multiple coincident signals. For example, the CRMP complex could sense mechanical forces in a manner analogous to the Piezo ion channels (Murthy *et al*, 2017), while the CLAMP complex may signal attachment to specific ligands on the host cell surface. Alternatively, rhoptry discharge factors may help define the site of rhoptry protein translocation.

The rhoptry discharge factors display varying conservation among alveolates. CRMPs and the Nd proteins (e.g. Nd9) were identified by their homology to factors involved in the mucocyst secretion machinery of ciliates, and show broad conservation across apicomplexans (Mageswaran *et al*, 2021; Aquilini *et al*, 2021; Sparvoli *et al*, 2022) By contrast, orthologs of MIC8 are limited to *T. gondii* and its closest relatives (Li *et al*, 2003), suggestive of more recent adaptations, perhaps mediating interactions with cell types uniquely encountered by these parasites (Boothroyd, 2009). SPATR displays incomplete conservation across the phylum— orthologs are found in *T. gondii, N. caninum*, and *Plasmodium* spp. (Li *et al*, 2003). Notably, *P. falciparum* SPATR fails to complement *T. gondii* SPATR, implying species-specific functional diversification (Huynh *et al*, 2014). While in *T. gondii* cell culture SPATR is dispensable, *Pf*SPATR is essential during blood stages in culture. The latter corroborates prior work, which found that inhibitory antibodies can block *P. falciparum* invasion of blood cells (Chattopadhyay *et al*, 2003). In *P. berghei, Pb*SPATR is required for blood stages and is likely required in sporozoites, suggesting the CLAMP complex is important for host cell invasion in multiple stages of the parasitic life cycle (Gupta *et al*, 2020; Costa *et al*, 2022). Though we were unable to make a conditional knockdown strain for *Pf*CLIP, the PlasmoGEM project finds that the CLIP homolog in *P. berghei* is also essential during blood stages (Bushell *et al*, 2017). Together, we predict a central function for CLAMP and CLIP, based on their essential phenotypes and conservation throughout the Apicomplexa, with SPATR adapting and modifying the core functions of the CLAMP complex to the individual requirements of certain species.

Although CLIP orthologs are highly divergent, representatives can be identified based on the conserved motif in all the parasite genomes examined. Although we did not identify CLIP homologs in gregarines or hemogregarines, they were represented in the cyst forming coccidia, monoxenic coccidia, piroplasms, hemosporidia, and cryptosporidia. While divergent in length and specific motif composition, these homologs shared at least one highly conserved motif at their C terminus, proximal to the single TM helix present in CLIP homologs—and the likely region interacting with CLAMP. In *T. gondii*, we identified two additional CLIP homologs: TGGT1_212275 and TGGT1_219742. While CLIP is regulated by the tachyzoite cell cycle in a manner consistent with other microneme proteins, TGGT1_212275 is upregulated in bradyzoites (Waldman *et al*, 2020). TGGT1_219742 is expressed at very low levels throughout the asexual lifecycle but appears to be upregulated in merozoites within the definitive host (Waldman *et al*, 2020; Hehl *et al*, 2015; Ramakrishnan *et al*, 2019). Intriguingly, this pattern of conservation is reminiscent of homologs of the moving junction components AMA1 and RON2, which are upregulated during different stages in the parasite life cycle (Lamarque *et al*, 2014).

Other cyst-forming coccidians have at least two CLIP homologs, whereas apicomplexans outside this clade typically possess a single homolog. As has been speculated for paralogs of AMA1 and RON2 (Lamarque *et al*, 2014), the diversification of CLIP paralogs may optimize function of the CLAMP complex during invasion in different tissue or host contexts.

This work describes a novel complex of three microneme proteins (CLAMP, CLIP, and SPATR) that mediates rhoptry content secretion in *T. gondii*. Patterns of conservation and essentiality in *Plasmodium* spp. lead us to conclude this complex likely performs analogous functions during invasion of host cells by other apicomplexan parasites. The discovery of the CLAMP complex will provide a foundation for future work aiming to identify the specific host and parasite interactions that trigger rhoptry secretion and the mechanisms underlying apicomplexan invasion.

## MATERIALS & METHODS

### *T. gondii* parasite maintenance and strain construction

*T. gondii* parasites from the strain RH and derived strains were cultured in human foreskin fibroblasts using Dulbecco’s Modified Eagle’s Medium (DMEM) supplemented with 3% fetal bovine serum or calf serum and 10 μg/ml gentamicin. Parasites and host cells were maintained at 37°C with 5% CO_2_. Pyrimethamine was used at 3 μM, xanthine (XA) was used at 50 μg/ml and mycophenolic acid (MPA) was used at 25 μg/ml.

CLAMP-cKD was constructed by transfecting pCLAMP-U1 (Sidik *et al*, 2016b) linearized with MfeI into Δku80/DiCre parasites (Pieperhoff *et al*, 2015) and selecting for integration with XA/MPA. Positive clones were isolated by limiting dilution and integration of the U1 construct was confirmed with immunofluorescence microscopy and western blotting using an anti-HA antibody (Biolegend cat. no. 901501). Parasite lysate was not boiled prior to western blotting, as we found that boiling reduced our ability to detect CLAMP. CLAMP depletion was induced by treating parasites with 50 nM rapamycin for 2 hours at least 24 hours prior to analysis.

All other strains were constructed using CRISPR-mediated homologous recombination, as described previously (Sidik *et al*, 2014). We added HA tags to CLIP in CLAMP-Ty (Sidik *et al*, 2016b) and Δku80/Tati (Meissner *et al*, 2002) parasites by transfecting these strains with pU6-Universal encoding a protospacer with the sequence CAGATGTGACTACCATTCGA along with a repair oligonucleotide made by duplexing the primers P1 and P2 (**Table S1**). To integrate the TetO7SAG4 promoter upstream of CLIP-HA in Δku80/Tati parasites, we amplified TetO7SAG4 from pDTS4 (van Dooren *et al*, 2008) using the primers P3 and P4. We transfected this PCR product into Δku80/Tati/CLIP-HA parasites along with pU6-Universal encoding a protospacer with the sequence GGTCAAGTACAGGAGTAATG and selected for integration. Positive clones were identified by PCR using the primers P5 and P6, and isolated using limiting dilution. A region spanning the junction the 3’ end of the Tet07SAG4 insert and the 5’ end of CLIP was amplified using the primers P7 and P8, and the junction was confirmed to be correct by Sanger sequencing using the primer P9. CLIP depletion was induced by treating parasites with 0.5-1 ug/ml anhydrous tetracycline for at least 24 hours prior to analysis.

CLAMP-Ty/SPATR-HA parasites were constructed by transfecting CLAMP-Ty parasites with pU6-Universal carrying a protospacer with the sequence CAAGCAGAGAACCATTCTGA along with a repair oligonucleotide made by duplexing primers P10 and P11. Positive clones were isolated using limiting dilution and confirmed using immunofluorescence microscopy.

### Generation of conditional knockdown *P. falciparum* lines

Conditional knockdown (cKD) *P. falciparum* parasites were generated for *Pf*CLAMP (PF3D7_1030200) and *Pf*SPATR (PF3D7_0212600) by fusing the coding sequences and noncoding RNA aptamer sequences in the 3’UTR, permitting translation regulation using the TetR-DOZI system (Ganesan *et al*, 2016; Nasamu *et al*, 2021). Gene editing was achieved by CRISPR/SpCas9 using the linear pSN054 vector containing cloning sites for the left homology region (LHR) and the right homology region (RHR) as well a gene-specific gRNA under control of the T7 promoter. Cloning into pSN054 was carried out following previously described procedures (Ganesan *et al*, 2016; Nasamu *et al*, 2021). The vector includes V5-2xHA epitope tags, a 10x tandem array of TetR aptamers upstream of an Hsp86 3’UTR, and a multicistronic cassette for expression of TetR-DOZI (regulation), blasticidin S-deaminase (selection marker) and a *Renilla* luciferase (RLuc) reporter. The LHR and re-coded regions were cloned in-frame with the tandem V5-2xHA tag such that CLAMP or SPATR were tagged at their C-terminus with the V5-2xHA tag post-editing, directly upstream of the Tet regulatory aptamer array embedded in the 3’UTR. All primer and synthetic fragment sequences were generated using the BioXP™ system and are included in Table S1. The final constructs were verified by restriction digestion and sequencing.

Transfection into Cas9 and T7 RNA polymerase-expressing NF54 parasites was carried out by pre-loading erythrocytes with the donor vector as previously described (Deitsch *et al*, 2001). Parasite cultures were maintained continuously in 500 nM anhydrotetracycline (ATc, Sigma-Aldrich cat. 37919) and drug selection with 2.5 μg/mL of Blasticidin S (RPI Corp B12150-0.1) was initiated four days after transfection. Cultures were monitored by Giemsa smears and RLuc measurements.

### *P. falciparum* growth assay

Assessment of proliferation rate upon *Pf*CLAMP and *Pf*SPATR knockdown used luminescence as a readout of growth. Synchronous ring-stage parasites cultured in varying concentrations of ATc (50, 3, 1, and 0 nM) were set up in triplicate in a 96-well U-bottom plate (Corning, cat. 62406-121). Dihydroartemisinin treatment (DHA; 500 nM) was used as a negative control for growth. Luminescence readings were taken at 0 and 72 h post-invasion using the Renilla-Glo^®^ Luciferase Assay System (Promega cat. E2750) and the GloMax^®^ Discover Multimode Microplate Reader (Promega). Luminescence values in the knockdown condition were normalized to maximum of ATc-treated (100% growth) and dihydroartemisinin (DHA) treated (500 nM, no growth) samples and results were visualized on a bar graph using GraphPad Prism (GraphPad Software).

### Immunofluorescence microscopy

To analyze CLIP and CLAMP’s localization, intracellular CLAMP-Ty and CLAMP-Ty/CLIP-HA parasites were fixed in 4% paraformaldehyde on ice for 20 minutes, then permeabilized with cold methanol for 2 minutes. Cells were stained with a 1:1000 dilution of mouse anti-Ty (Bastin *et al*, 1996), a 1:1000 dilution of mouse anti-MIC2 (Achbarou *et al*, 1991), or a 1:1000 dilution of mouse anti-HA (Biolegends cat. No. 901501) and Hoechst (Santa Cruz cat. no. 394039, 2000-fold dilution).

### Host cell adhesion assay

DiCre and DiCre/CLAMP-U1-HA parasites that switch from expressing killer red to YFP in response to rapamycin (Sidik *et al*, 2016b) were treated with 1 μM mycalolide B in Ringer’s (155 mM NaCl, 3 mM KCl, 2 mM CaCl_2_, 1 mM MgCl_2_, 3 mM NaH_2_PO_4_, 10 mM HEPES, 10 mM Glucose, 1% fetal bovine serum) for 30 minutes at room temperature, then resuspended at 2 x 10^6^ parasites/ml in Ringer’s solution lacking mycalolide B. These parasites were perfused through microfluidic channels (Ibidi cat. no. 80196) seeded with HFF cells at a rate of 20 μl/min for 5 minutes, then at a rate of 50 μl/min while imaging at 10X magnification every 2 seconds for 5 minutes. Videos were analyzed with a custom script that identified attached parasites, which were defined as parasites whose centers of mass did not move more than 5 pixels between any 2 consecutive frames for at least 20 seconds. The number of attached parasites was normalized to the average number of parasites per frame of a given video to account for small differences in parasite concentrations. When parasites were treated with rapamycin, only YFP-expressing parasites were analyzed. HFF cells were pre-treated with 10 mg/ml heparin in Ringer’s for approximately 1 hour when indicated.

### Coupled analysis of the total proteome and ESA fractions

At the time of infection, ten 15 cm plates of HFF cells were infected with the CLAMP-HA cKD strain were treated with 50 nM rapamycin or vehicle control, such that there were six 15 cm tissue culture plates for rapa-treated parasites and 4 for the vehicle treated condition. After 2 hours, the rapamycin was removed with 3x PBS washes and a D3C wash. For the CLIP-HA cKD strain, HFF cells were infected in presence of 0.5 μg/μL anhydrotetracycline or a vehicle control, with five 15 cm cell culture dishes infected for each treatment condition. Forty-eight hours later, extracellular parasites were filtered using 3 μm membranes to remove host cell debris. Parasites were pelleted at 1000 g for 5 minutes at 4°C, washed in DMEM with 0% serum (D0), centrifuged again and resuspended in 130 uL D0 for each plate of parasites. For the ESA fraction experiments, 75 μL of parasites for each condition were then pretreated with 75 μL of 300 μM BAPTA-AM in D0 or a DMSO vehicle control in 1.5 mL microcentrifuge tubes, and incubated in a 37°C water bath for 15 minutes. For the ESA fractions, 75 μL of 750 μM zaprinast or a DMSO vehicle control was added to each tube, then tubes were further incubated in a 37°C water bath for 30 min. The parasites were then spun down (4°C, 1000 g, 5 minutes) and the ESA-containing supernatant was pipetted into a protein LoBind tube. The supernatant was then spun down and the supernatant removed a second time to a new protein LoBind tube to ensure removal of all parasites.

During the zaprinast incubation, the remaining parasites not used for generation of the ESAs were spun down and resuspended in Ringer’s lysis buffer lysed down, then lysed in Ringer’s containing 0.85% NP-40 (155 mM NaCl, 3 mM KCl, 2 mM CaCl_2_, 1 mM MgCl_2_, 3 mM NaH_2_PO_4_, 10 mM HEPES, 10 mM Glucose, 0.8% Igepal CA-630, Halt protease inhibitors (Life Technologies cat. no. 87786)).

For MS analysis, parasite lysates were treated with 25 units of benzonase nuclease for 10 minutes at room temperature, then both lysate and ESA samples were desalted and cleared of detergents using hydrophilic and hydrophobic SP3 beads (GE Healthcare cat. no. 45152105050250 and 65152105050250) as previously described (Hughes *et al*, 2019). Proteins were resuspended in 50 mM triethylammonium bicarbonate (TEAB, Sigma cat. no. T7408), reduced for 1 hour at 55°C using 10 mM tris(2-carboxyethyl)phosphine (TCEP, Pierce cat. no. 20490), alkylated for 1 h at 24°C using 25 mM iodoacetamide (CLAMP-HA cKD experiments, Sigma cat. no. I1149) or methyl methanethiosulfonate (CLIP-HA cKD experiments, Pierce cat. no. 23011), then digested overnight using sequencing grade modified trypsin (Promega cat. no. V5113). Peptides were incubated for 1 h at 22°C in the presence of a 2:1 mass ratio of TMT10plex reagent (Life Technologies cat. no. 90111) to peptides, and excess TMT10plex reagent was quenched using addition of hydroxylamine to 0.3%. ESA samples were combined and desalted using a Sep-Pak Light cartridge (Waters cat. no. WAT023501) while lysate samples were combined, then desalted and fractionated using a reversed-phase fractionation kit (Pierce cat. no. 84868). Peptides were analyzed using a Thermo Q Exactive HF-X Hybrid Quadrupole-Orbitrap mass spectrometer, and downstream analysis was performed using Proteome Discoverer Version 2.4.

For immunoblot analysis of ESA fractions of CLIP-HA cKD, the indicated ESA supernatant was boiled in SDS-PAGE sample buffer and 3 μg of each sample was run with a 4-15% TGX gradient gel (Bio-Rad cat. no. 4561086), then transferred to a nitrocellulose membrane. The membrane was immunoblotted with anti-CLIP-N and anti-MIC2 (Achbarou *et al*, 1991)

### Anti-CLIP-N antibody generation

*E. coli* codon optimized CLIP coding sequence (TGGT1_212270) for residues 1-187 was Gibson-cloned into pVP57K, resulting in a His-MBP-TEV-CLIP-1_187 expression construct.

For protein expression pVP57K-CLIP-1_187 was transformed into BL21 Star (DE3) *E. coli* (Invitrogen cat. no. K10101), then grown with shaking at 37°C in LB media to an optical density at 600 nm (OD600) of between 0.6 and 0.8, then cooled quickly on ice and induced with 0.5 mM Isopropyl β-D-thiogalactoside (IPTG, Sigma cat. no. I6758) and allowed to incubate with shaking overnight at 16°C. The cells were pelleted by centrifugation (3000 g, 4°C) and resuspended in a 50 mL tube with 40 mL of binding buffer (250 mM NaCl, 50 mM Tris 7.5, 10 mM imidazole, 2.5% glycerol), and frozen in liquid nitrogen. For purification, the cells were thawed on ice, 1 mM phenylmethylsulfonyl fluoride (PMSF, Sigma cat. no. 10837091001) was added, and the sample was homogenized with a microfluidizer (Microfluidics M-110L). The soluble fraction was separated from debris by centrifugation (9000 g, 4°C) and incubated for 60 minutes with Ni-NTA beads (Qiagen, cat. no. 30230) pre-equilibrated in binding buffer. The resin was captured on a gravity flow column and washed with >100 mL wash buffer (binding buffer with 35 mM imidazole). His-MBP-TEV-CLIP-1_187 was eluted from the resin by 2 mL volumes of elution buffer (binding buffer with 300 mM imidazole) until elution was complete. The protein was then dialyzed overnight to remove the imidazole (250 mM NaCl, 50 mM Tris pH7.5, 2.5% glycerol, 0.25 M EDTA, 0.5 mM DTT), concentrated using a 10,000 MWCO spin concentrator (Amicon) to >2 mg/mL, the particulates were removed with a 0.22 μm spin filter (Corning cat. no. 8160) and the protein was aliquoted into ~200 ug samples and stored at −80°C.

Custom rabbit anti-CLIP-N antiserum was made by Covance Laboratories Inc. Briefly, 250 μg of His-MBP-TEV-CLIP-1_187 protein emulsified with Freund’s Complete Adjuvant was used for the primary immunization, followed by three boosts of 125 μg antigen with Freund’s Incomplete Adjuvant at periods of 21 days starting from the initial immunization (day 0). The final termination bleed is referred to as anti-CLIP-N and reactivity towards endogenous CLIP confirmed by immunoblot.

### Electron microscopy

Extracellular parasites were treated with 1 μM mycalolide B or 1 μM cytochalasin D in invasion media (sodium bicarbonate-free DMEM containing 20 mM HEPES and 1 % fetal bovine serum, pH 7.4) for 30 minutes at room temperature, then combined with K562 cells at an MOI of ~10. Parasites and host cells were incubated at 37°C for 15-30 minutes, then fixed for 45 minutes on ice with 2% OsO_4_ and 1% glutaraldehyde in 0.5X phosphate buffer (1X phosphate buffer consists of 38.4 mM K_2_HPO_4_, 161.6 mM KH_2_PO_4_, pH 6.2). Samples were rinsed in cold dH_2_O, then stained en bloc staining with 1 % aqueous uranyl acetate (Ted Pella Inc.) at 4 °C for 3 h. Samples were rinsed, then dehydrated in a graded series of ethanol and embedded in Eponate 12 resin (Ted Pella Inc.). Ninety-five nanometer sections were cut with a Leica Ultracut UCT ultramicrotome (Leica Microsystems Inc., Bannockburn, IL), stained with uranyl acetate and lead citrate, and imaged on a JEOL 1200 EX transmission electron microscope (JEOL USA Inc.) equipped with an AMT 8 megapixel digital camera and AMT Image Capture Engine V602 software (Advanced Microscopy Techniques).

### Evacuole and pSTAT6 assays

To visualize evacuoles, extracellular parasites were suspended in 0.5 μM cytochalasin D in endo buffer (44.7 mM K_2_SO_4_, 10 mM Mg_2_SO_4_, 106 mM sucrose, 5 mM glucose, 20mM Tris pH 8.2, 0.35% (wt/vol) BSA), then applied to monolayers on HFF cells grown on coverslips.

Parasites and HFF cells were incubated at 37°C with 5% CO_2_ for 15 minutes to allow the parasites to adhere to the host cells, then the media was gently replaced with 1 μM cytochalasin D in HHE (HBSS (Sigma cat. no. H2387) containing 0.1 mM EGTA and 10 mM HEPES) and parasites and host cells were incubated at 37°C with 5% CO_2_ for an additional 20 minutes. Cells were fixed in cold methanol for 10 minutes, then stained with anti-ROP1 (Bradley & Boothroyd, 1999), anti-CDPK1 (Waldman *et al*, 2020), and Hoechst (Santa Cruz cat. no. 394039).

To visualize parasite-induced nuclear accumulation of host pSTAT6, parasites were pretreated with 1 μM mycalolide B (VWR cat. no. 89165-002) in invasion media (sodium bicarbonate-free DMEM containing 20 mM HEPES, pH 7.4) supplemented with 1% fetal bovine serum for 30 minutes at room temperature, then washed 3 times with PBS. Parasites concentrations were then normalized to 1 x 10^6^ parasites per ml in DMEM containing 3% serum and 10 μg/ml gentamicin, and 200 μl of parasites were applied to each well of HFF cells grown in 96-well plates with transparent bottoms (Perkin Elmer cat. no. 6055300). Plates were centrifuged at 290 g for 5 minutes, then incubated at 37°C with 5% CO_2_ for 4 hours. Cells were fixed in 4% formaldehyde on ice for 20 minutes, then permeabilized in PBS containing 0.25% Triton X-100, 5% fetal bovine serum and 1% normal goat serum for 8 minutes at room temperature. Cells were stained using anti-pSTAT6 (Cell Signalling cat. no. 9361S or Abcam cat. no. ab188080), anti-CDPK1 (Waldman *et al*, 2020), and Hoechst (Santa Cruz cat. no. 394039). Cells were imaged using a Cytation 3 high-content plate reader (BioTek), and pSTAT6-positive host cell nuclei were counted using a custom image analysis script.

### Immunoprecipitations

For anti-Ty immunoprecipitations, Ty antibody (Bastin *et al*, 1996) was conjugated to agarose beads carrying protein G (Life Technologies cat. no. 22851) using 22 mM DMP, then washed in 0.2 M sodium borate pH 9.0 and 0.2 M ethanolamine, 0.2M NaCl pH 8.5. Beads were washed in 100 mM glycine pH 2.5 followed by neutralization buffer (50 mM HEPES pH 7.4, 1 mM EGTA, 1 mM MgCl_2_, 300 mM KCl, 10% glycerol) to remove excess antibody prior to use. For reciprocal immunoprecipitations, pre-conjugated magnetic anti-HA beads were used (Pierce 88836).

For all IPs, approximately 3-5 x 10^8^ Parasites were lysed in lysis buffer (150 mM NaCl, 20 mM Tris pH 8.0, 0.1% SDS, 1% Triton X-100, Halt protease inhibitors (Life Technologies cat. no. 87786)), then cell debris was removed by centrifugation. In the case of anti-Ty IPs, lysates were additionally precleared by incubating with Protein G slurry for ~1 h at 4°C. Antibody-conjugated Ty or HA beads (Pierce 88836) that had been pre-washed with lysis buffer were then incubated with lysates for for ~1h at 4°C. Beads were washed in lysis buffer, then conjugated proteins were eluted using 200 mM glycine pH 2.5, which was neutralized with an equal volume of 1 M Tris pH 8.5 prior to SDS-PAGE or MS analysis.

### CLIP phylogenetic analysis

ToxoDB (Gajria *et al*, 2008), PlasmoDB (Aurrecoechea *et al*, 2009), and CryptoDB (Heiges *et al*, 2006) were searched for CLIP homologs. Clustal Omega (Madeira *et al*, 2019) was used to align these homologs in the PHYLIP file format, which was then used with the seqboot, proml, and consense programs from the PHYLIP phylogenetics package (Felsenstein, 1989). Bootstrapping was performed 100 times, and the input order was randomized twice when using proml. The Jones-Taylor-Thornton probability model was used. Output from consensus was used to draw trees in Figtree (http://tree.bio.ed.ac.uk/software/figtree/). Motifs that are shared by CLIP homologs were identified using the MEME suite for motif-based sequence analysis (Bailey *et al*, 2009).

### Membrane association assays

Approximately 1 x 10^8^ CLAMP-Ty or CLAMP-Ty/CLIP-HA parasites were passed through a 3 μm filter to remove host cell debris, then split into two samples. Parasites were pelleted and resuspended in IC buffer (1.37 M KCl, 50 mM NaCl, 200 mM HEPES, 100 mM MgCl2, Halt protease inhibitors) with or without 2% Triton X-100. Samples were cycled between liquid nitrogen and a 37°C water bath three times, then disrupted 5 times with a dounce homogenizer (VWR cat. no. 62400-595). Following mechanical disruption, samples were centrifuged at 100,000 RPM (~400,000 g) for 1 h. Supernatants were collected, and pellets were resuspended in IC buffer containing 2% Triton X-100. Samples were immunoblotted using anti-Ty (Bastin *et al*, 1996), anti-Actin (Dobrowolski *et al*, 1997), anti-SAG1 (Burg *et al*, 1988), anti-HA (Biolegend cat. no. 901501), and anti-Aldolase (Starnes *et al*, 2009).

### Proteinase K protection assays

Approximately 2 x 10^8^ CLAMP-Ty parasites were passed through a 3 μm filter to remove host cell debris, then resuspended in cold SoTE (0.6 M sorbitol, 20 mM Tris-HCl pH 7.5, 2 mM EDTA, pH 7.5). The parasites were split into 10 tubes, and digitonin was added at final concentrations ranging from 0.001 to 0.008%. Results shown are from samples treated with 0.008% digitonin. Parasites were incubated on ice for 3 minutes, then spun at 4°C and 600 x *g* for 10 minutes. Parasites were then resuspended in SoTE with or without 100 μM proteinase K and incubated on ice for 30 minutes. Proteins were precipitated using trichloroacetic acid and immunoblotted using anti-Ty (Bastin *et al*, 1996) and anti-MIC2 (Achbarou *et al*, 1991).

### Plaque assays

Five hundred CLIP-cKD or parental strain parasites were applied to monolayers of HFF cells in the presence of 1 μg/ml anhydrous tetracycline or vehicle control. Cells were incubated at 37°C with 5% CO_2_ for 8 days, then washed in PBS and fixed with 100% ethanol. Monolayers were stained with crystal violet solution (2% crystal violet, 0.8% ammonium oxalate, 20% ethanol), washed with water, and air-dried.

### Invasion assays

Gene depletion was induced using either 50 nM rapamycin for DiCre/CLAMP-HA-U1 (CLAMP-HA cKD) or 0.5 μg/ml anhydrous tetracycline for TATi/TetO7-CLIP-HA (CLIP-HA cKD) two days prior to assaying invasion. 2 x 10^5^ parasites were applied to each well of a clear-bottomed 96-well plate (Perkin Elmer cat. no. 6055300) seeded with HFF cells. Three wells were used for each strain. Plates were centrifuged at 290 x *g* for 5 minutes, then incubated at 37°C for 10 minutes. Cells were fixed in 4% formaldehyde on ice for 20 minutes, then extracellular parasites were stained with anti-SAG1 (Burg *et al*, 1988) prior to permeabilization. Cells were permeabilized with 0.25% Triton X-100 in PBS containing 5% fetal bovine serum and 1% normal goat serum, then all parasites were stained with anti-CDPK1 (Waldman *et al*, 2020) and host cell nuclei were stained with Hoechst (Santa Cruz cat. no. 394039). Assays were imaged using a Cytation 3 high-content plate reader (BioTek). Intracellular parasites, extracellular parasites, and host cell nuclei were identified using a custom image analysis script, and the number of intracellular parasites was normalized to the number of host cells.

---

**Figure S1.**
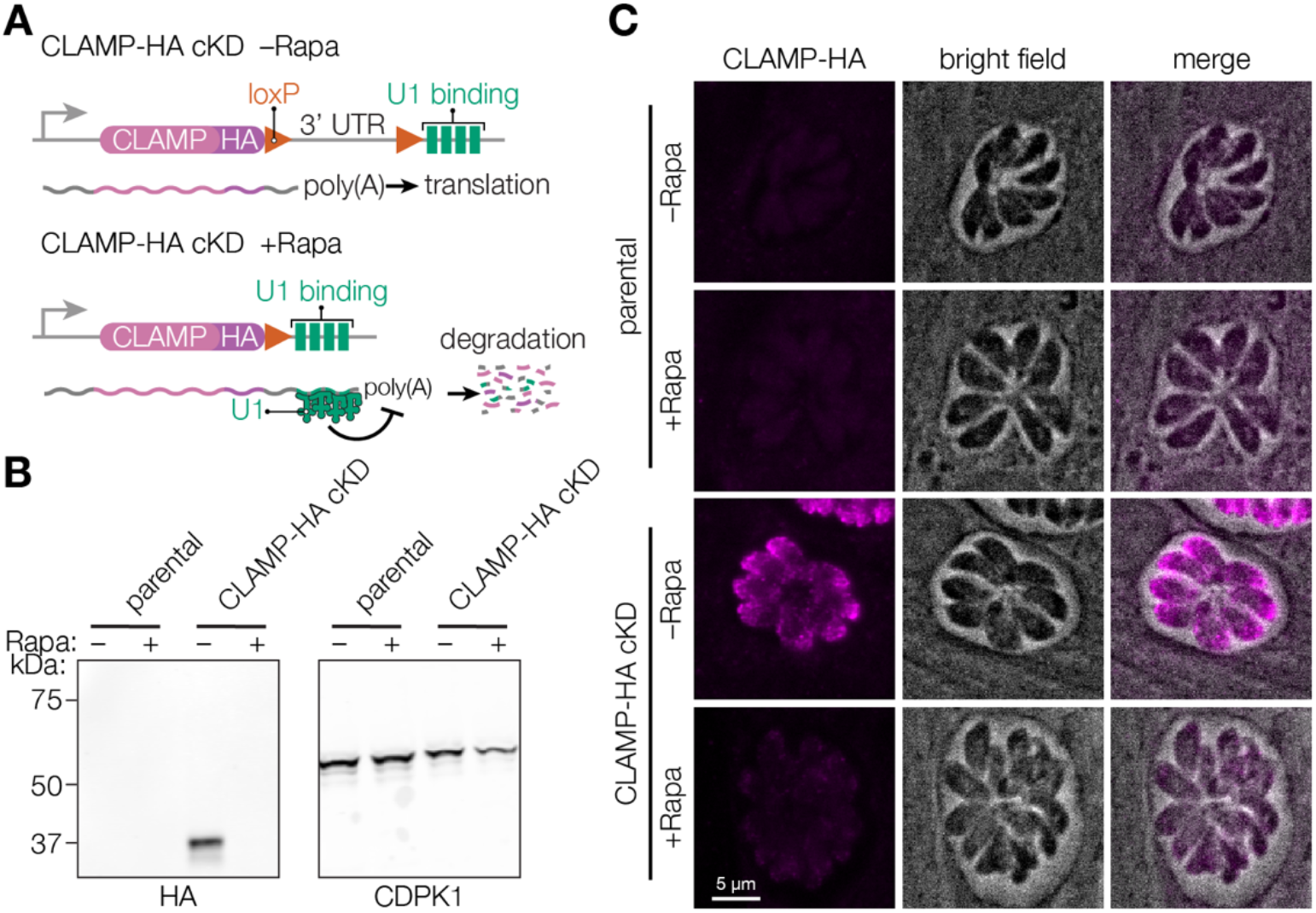
Generation of a non-fluorescent CLAMP conditional knockdown in *T. gondii* strain. **A**. Diagram of the CLAMP-HA cKD scheme. Induction of DiCre dimerization by rapamycin (Rapa) treatment leads to excision of the 3’UTR between the stop codon and an array of U1 binding sites, leading to U1-mediated degradation of the mRNA transcript. **B–C**. Knockdown of CLAMP-HA expression was assessed following a 2 h treatment with rapamycin. Samples were analyzed 48 h after the treatment by anti-HA immunoblotting (B), or after 24 h after the treatment by anti-HA immunofluorescence (C).

**Figure S2.**
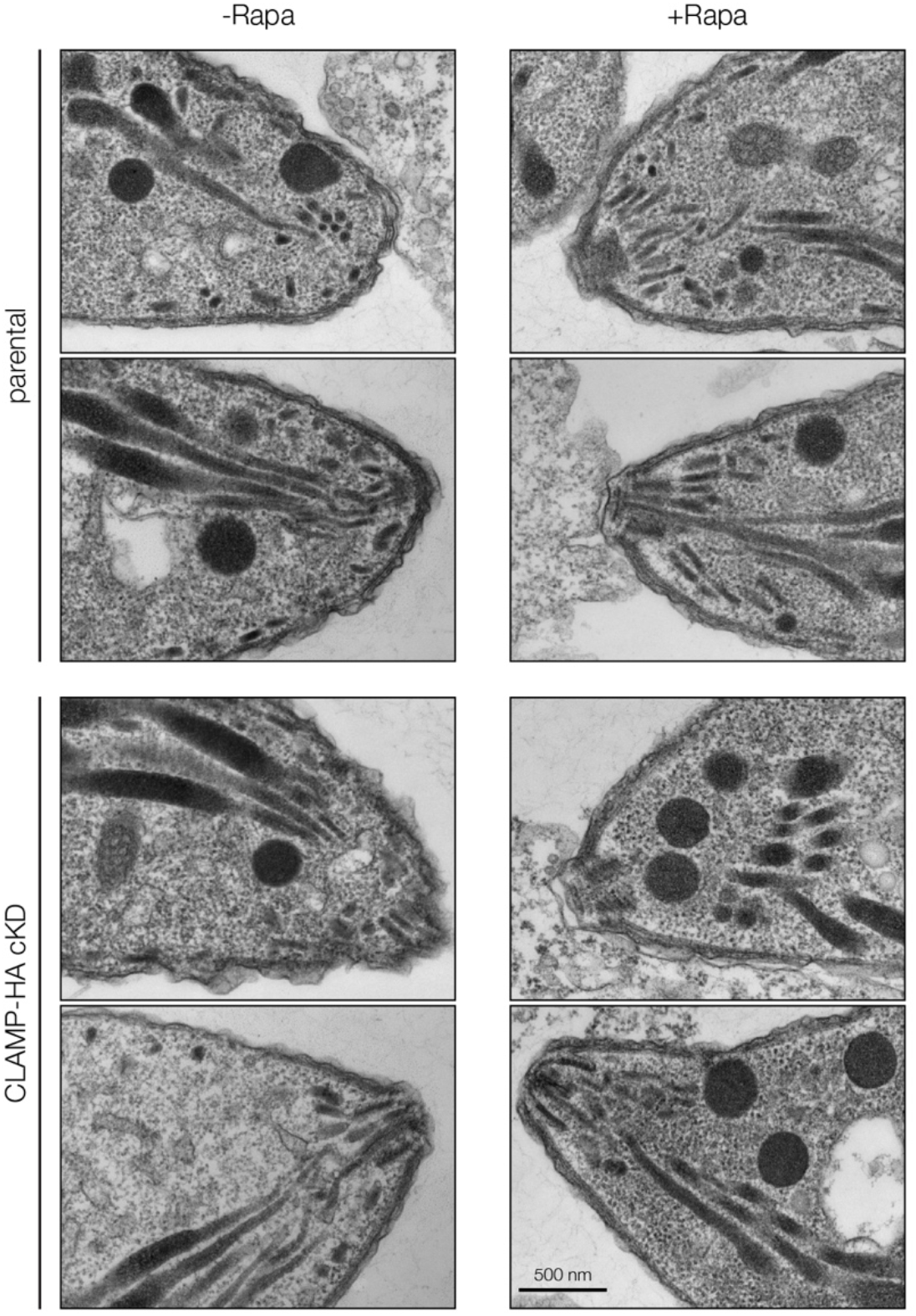
CLAMP knockdown does not alter apical-complex morphology in *T. gondii*. Additional images of CLAMP-HA cKD and parental parasites prepared for electron microscopy 48 h after a 2 h treatment with 50 nM rapamycin (+Rapa) or a DMSO vehicle control (−Rapa).

**Figure S3.**
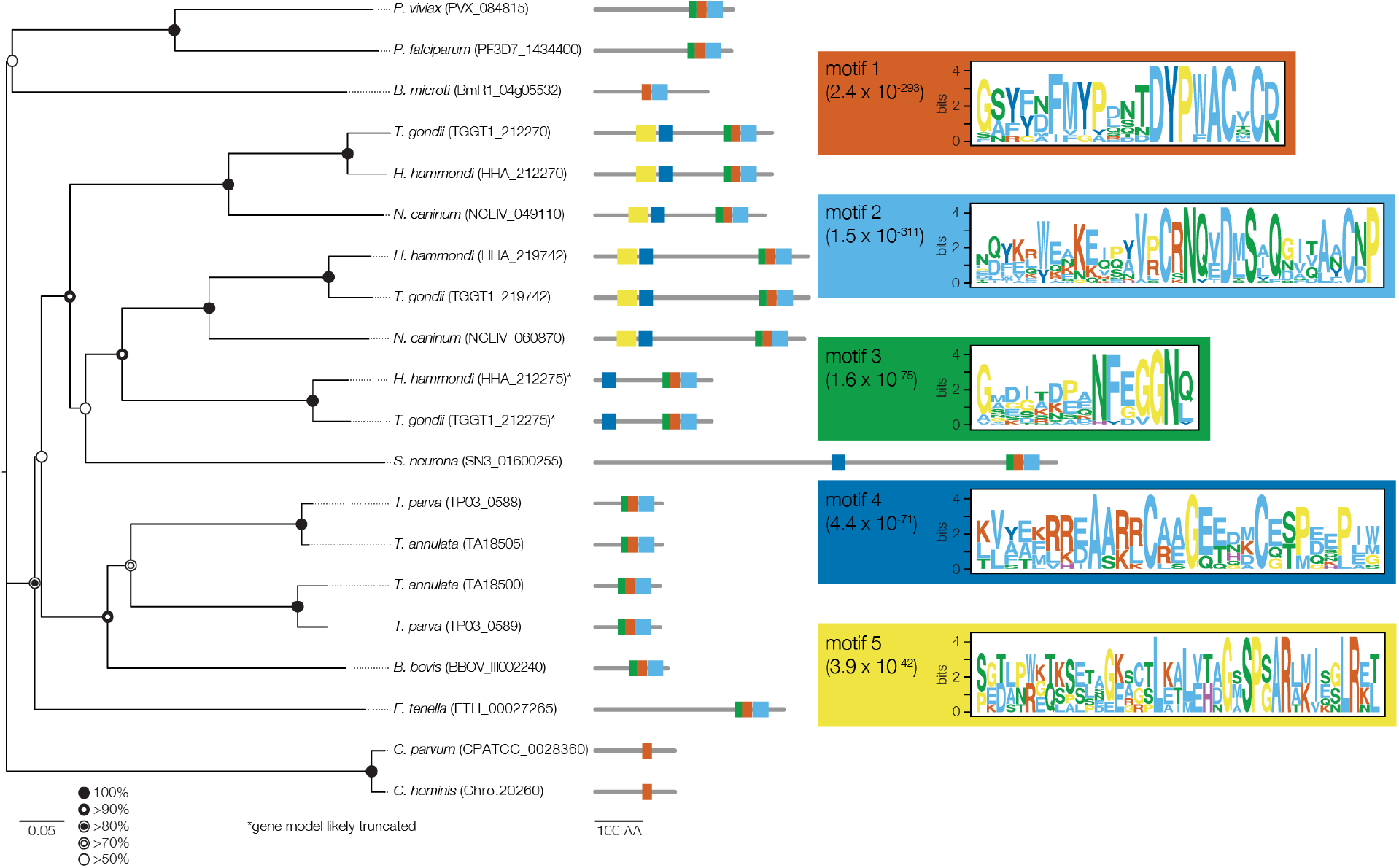
Extended analysis of CLIP homologs. Neighbor-joining phylogenetic tree of CLIP homologs. Bootstrap values for 1,000 trials are displayed. The logo plots for the top five motifs identified using MEME analysis are shown in boxes colored to match the domains on the protein models. E-value for each motif within the set of 20 CLIP homologs is indicated in parentheses. Sequence analysis suggests some of the models are incomplete (*), and a *N. caninum* sequence syntenic with TGGT1_212275 could be detected, but is not annotated in the current version of the genome (ToxoDB release 60). Abbreviated species names are provided for *Toxoplasma gondii, Plasmodium vivax, Plasmodium falciparum, Theileria parva, Theileria annulata, Babesia bovis, Babesia microti, Neospora caninum, Sarcocystis neurona, Hammondia hammondi*, *Eimeria tenella*, and *Cryptosporidium parvum*.

**Figure S4.**
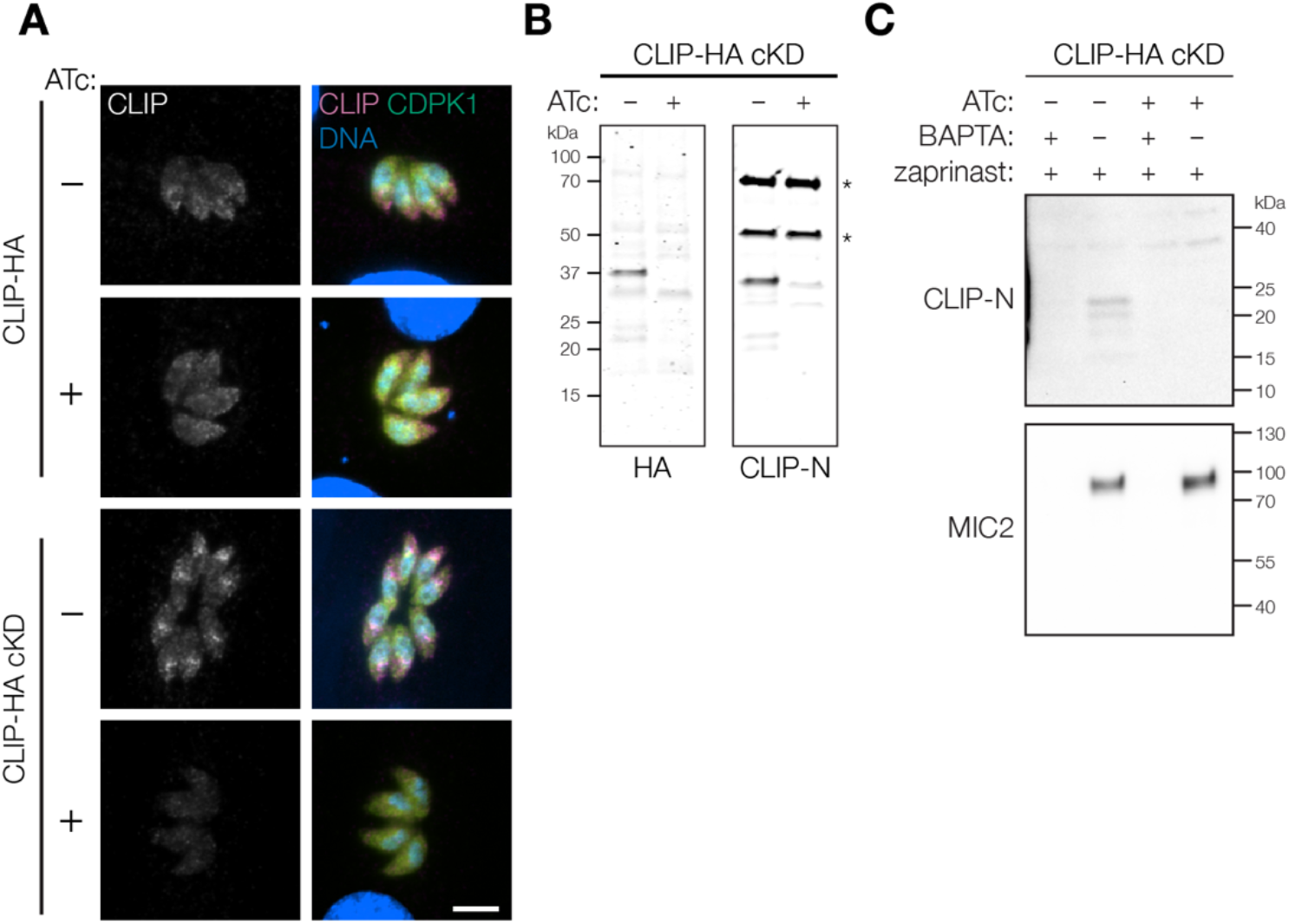
Conditional CLIP knockdown disrupts microneme localization and blocks detection of CLIP in the extracellular secreted antigen fraction. **A.** Treatment with anhydrotetracycline (ATc) leads to loss of CLIP-HA signal in the CLIP-HA cKD strain, as visualized by anti-HA immunofluorescence. **B**. To improve detection of the N-terminal region of CLIP without the requirement of an internal epitope, an antibody was generated against the N-terminal 187 residues of CLIP (CLIP-N). Probing the CLIP-HA cKD strain after a 2-day treatment with ATc or a vehicle control demonstrates the antibody sensitively detects CLIP. Non-specific bands detected by the anti-CLIP-N serum are marked with an asterisk (*). **C**. Probing the MS-ESA samples from Figure 4I–J with anti-CLIP-N confirms that a form of cleaved CLIP is found in the ESA fraction, as is the microneme protein MIC2. Knockdown of CLIP with ATc leads to loss of CLIP in the ESA fraction. As expected, microneme secretion is inhibited by BAPTA-AM and induced by zaprinast treatment.

## SUPPLEMENTAL MATERIAL

**Supplemental Table 1.** Primers and dsDNA fragments used to generate the parasite strains used in this study.

**Supplemental Table 2.** Mass spectrometry data for CLAMP-Ty immunoprecipitation experiments.

**Supplemental Table 3.** Mass spectrometry data for MS-ESA and whole proteome analysis for CLAMP cKD and CLIP cKD experiments.

**Supplemental Video 1.** *Toxoplasma* adhesion of parental control strain with and without heparin treatment.

**Supplemental Video 2.** Adhesion of CLAMP cKD with either vehicle or rapamycin treatment.

## ACKNOWLEDGEMENTS

We would like to thank L. David Sibley and Peter Bradley for the MIC2, SAG1, and ROP1 antibodies, Wandy Beatty and the Molecular Microbiology Imaging facility at the Washington University in St. Louis for assistance with the electron microscopy, Sarvesh Varma for assistance with microfluidics, Markus Meissner for the MIC8 cKD strain, and Gary Ward for the AMA1 cKD strain. We thank all members of the VEuPathDB team, which provided invaluable resources for the preparation of this manuscript. This project was supported by a grant from the National Institutes of Health to S.L. (R01AI144369), a Bill & Melinda Gates Foundation grant to J.C.N. (INV-026505), and a Long-term Fellowship from the Human Frontiers Science Program (LT000890/2021-L) to D.V.

## AUTHOR CONTRIBUTIONS

S.M.S., D.V. and S.L. designed the overall study. S.M.S, D.V., Y.A., C.F.A.P., and L.C.G. were responsible for performing experimental investigations. Y.A. performed MIC2 and AMA1 pSTAT6 assays. C.F.A.P., L.C.G., and J.C.N. were responsible for *P. falciparum* strain conceptualization, strain generation, validation, and growth assays. *Tg*SPATR knockout and complement strain generation and validation was performed by M.H. and V.B.C., as well as generation of anti-*Tg*SPATR polyclonal sera. S.M.S. and D.V. performed remaining experiments and analysis. S.M.S., D.V. and S.L. wrote the manuscript, and all other authors reviewed and approved the manuscript.

